# A highly dynamic active state for transducin-bound phosphodiesterase-6 in vertebrate phototransduction

**DOI:** 10.64898/2026.04.01.715611

**Authors:** Jessica N. Holechek, Julia Y. Shang, Tufa E. Assafa, Brian R. Crane, Richard A. Cerione

## Abstract

In vertebrate phototransduction, the G protein-coupled receptor rhodopsin activates the α-subunit of transducin (Gα_T_), which, upon binding the γ-subunits of phosphodiesterase-6 (PDE6), stimulates cGMP hydrolysis. We reported a cryoEM structure for a complex containing two constitutively active Gα_T_ (Gα_T_*) subunits coupled by a bivalent antibody bound to PDE6 that demonstrated a striking displacement of both PDEγ subunits from the PDEα/PDEβ catalytic sites and suggested an alternating-site mechanism for PDE6 activation. Here, we use site-directed spin labeling (SDSL) and double electron-electron resonance spectroscopy (DEER) to probe PDE6 conformational changes upon Gα_T_* binding. PDEγ spin-labelled on Cys68 and Ile64Cys demonstrate that PDEγ have highly flexible C-termini that transiently bind to the PDEα/PDEβ heterodimer. Binding of Gα_T_* to PDE6 with the inhibitor udenafil occupying its catalytic sites alters the positions of the PDEγ subunits in agreement with the changes shown in the cryoEM structure for this complex, whereas coupling the Gα_T_* subunits to the bivalent antibody does not affect the DEER distributions observed for PDE6 bound to Gα_T_*. However, binding of the slow hydrolyzing 8-Br-cGMP substrate in the presence of Gα_T_* causes a dramatic increase in the separation and spread of the spin-labelled PDEγ subunits, thereby revealing a previously unobserved conformation of PDE6 associated with catalysis, which is further supported by small angle X-ray scattering (SAXS) analysis. These studies indicate that whereas inhibitors trap Gα_T_*-PDE6 complexes in an inactive state as represented by the cryoEM structure, the binding of both substrate and Gα_T_* produces a dynamic active state consistent with an alternating-site mechanism.

## Introduction

The molecular machinery of rod photoreceptors engenders a highly amplified signaling cascade necessary for vision in low light intensities. This visual transduction pathway begins with the photoexcitation of the prototype class A G protein receptor (GPCR) rhodopsin (1). The activation of rhodopsin leads to conformational changes in the receptor that enables the binding and activation of its G protein partner transducin. The alpha subunit of transducin (Gα_T_) then interacts with the cGMP phosphodiesterase-6 (PDE6), which catalyzes the hydrolysis of cGMP to GMP (2). This phototransduction sensory response pathway is one of the most highly amplified signaling systems in biology. Photoexcited rhodopsin catalyzes the activation of as many as 10^2^ transducin molecules per cycle, whereas Gα_T_ activation of PDE6 leads to the hydrolysis of 10^3^ cGMP molecules per second (3). The resulting rapid decrease in cGMP concentration results in the closing of cyclic nucleotide-gated ion channels and subsequent hyperpolarization of the rod cells, leading to the physiological response of vision in dim light (4). Dysregulation of PDE6 and its control of cGMP levels has been implicated in retinitis pigmentosa and congenital stationary night blindness, and ultimately progressive vision loss (5, 6).

PDE6 is the central effector enzyme within the phototransduction pathway in retinal rods and consists of two similar but non-identical α and β subunits (PDEα; 99 kDa and PDEβ; 98 kDa). Each subunit contains a C-terminal catalytic domain and two N-terminal tandem GAFa and GAFb regulatory domains. Two critical autoinhibitory PDEγ subunits (∼10kDa) make up the heterotetrameric PDE6 complex. The two PDEγ subunits interact with both the catalytic and regulatory GAF domains of the PDEα/PDEβ heterodimer to regulate the cGMP hydrolytic activity of the enzyme, preventing basal cGMP hydrolysis in the absence of Gα_T_ (7). The autoinhibitory subunits allow for the extraordinarily high signal-to-noise ratio required for vision in dim light. Structural studies suggest that the binding of activated Gα_T_ subunits to PDE6 causes striking changes in the positions of the PDEγ subunits relative to the PDEα/PDEβ catalytic sites that are necessary for the stimulation of cGMP hydrolysis (8, 9). Whereas this phototransduction pathway has been long studied and the activation of transducin by the GPCR rhodopsin has been probed in detail, less is known about the structural and mechanistic basis underlying how Gα_T_ elicits such a large stimulation of PDE6 catalytic activity through binding to the PDEγ subunits.

The PDE6 heterodimer has been suggested to have two distinct binding sites for the PDEγ subunits (10, 11), although there have been suggestions that cGMP hydrolysis does not occur simultaneously, nor independently at the two catalytic sites on PDE6. For example, the binding of PDEγ to only one of its two sites on PDE6 was shown to be sufficient to inhibit enzyme activity (12). Additionally, maximal cGMP hydrolysis by PDE6 treated with saturating amounts of Gα_T_ only reaches 50% of the activity measured when both PDEγ subunits are removed from PDE6 by limited proteolysis (13), indicating that the two sites are not activated simultaneously and pointing towards an alternating-site mechanism. The high-affinity binding of PDEγ is mediated primarily through residues within three regions, namely the polycationic region (residues 27-38 on PDEγ), the glycine-rich region (residues 52-54), and the C-terminal end of PDEγ (residues 73-87) (14, 15). The binding of the C-terminal region of PDEγ was originally thought to inhibit PDE6 by blocking substrate access to the active site such that displacement of this C-terminal region by Gα_T_ would then stimulate PDE6 catalytic activity (15).

However, recent cryoEM studies indicate that even in the absence of Gα_T_, the competitive inhibitors udenafil and vardenafil displace the PDEγ C-termini and can access both active sites, thereby suggesting that the PDEγ C-termini also do not directly block the binding of substrates (16). Furthermore, active site occupation has been shown to be critical for the binding of Gα_T_ to PDE6 as Gα_T_ was only visible in 2D classifications of the complex that had been previously treated with the competitive inhibitor vardenafil (16). Allosteric inhibition independent of the C-terminal end of PDEγ that requires Gα_T_ to relieve inhibitory constraints in other regions of PDEγ has also been suggested (17).

Gα_T_ engages PDEγ through key residues in the glycine-rich and C-terminal regions (15). Biochemical studies have shown that two GTP-bound Gα_T_ subunits are required to achieve maximal activation of PDE6 (18), and that the two Gα_T_ subunits bind with different affinities to the two PDEγ subunits on PDE6 (19). The only current high-resolution cryoEM structure of the PDE6-transducin complex depicts two Gα_T_ subunits displacing the C-terminal ends of both PDEγ subunits from the catalytic domains on PDE6 and binding to the GAFb domains (9). Principal component analysis (PCA) of the PDE6-transducin cryoEM structure provided indications of alternating motions between the two Gα_T_ subunits (9). Isolation of this complex required occupation of the active site with vardenafil and the use of a bivalent antibody (Rho1D4) that recognizes the C-terminal ends of an engineered constitutively active Gα_T_, denoted Gα_T_*. The Gα_T_* subunit contains two residue substitutions (Q200L and R174C) that prevent GTP hydrolysis, and a C-terminal Rho1D4 epitope (9). Whereas this structure was the first to provide high resolution structural information on the complex, several unexpected features were resolved: striking reorientation of the PDEγ subunits away from their position in the apo-state, an upside-down conformation of Gα_T_* relative to the presumed location of the plasma membrane, and a symmetric doubly-bound state (9). These findings raised concerns that the antibody might influence the observed conformations, prompting us to investigate this complex using alternative techniques. In structural studies of this complex formed in the absence of the bivalent antibody, 2D classifications and 3D reconstructions showed weak density for 2 Gα_T_ subunits bound to the PDEα/PDEβ heterodimer with PDEγ not being resolved (16). The “apo” structure of PDE6 in the absence of Gα_T_ was also resolved at high resolution (PDB: 8ulg). It reveals full length PDEγ subunits that includes density for a previously undiscerned region of residues 49-71. Surprisingly, without the Gα_T_ subunits bound, there is little change in the orientation or conformation of the catalytic domains relative to the complex.

Although these cryoEM structures have contributed greatly to our understanding of this complex interaction, information on the positioning and structural fluidity of the PDEγ subunits has been difficult to obtain owing to their flexible nature. Paired with traditional structural techniques, double electron-electron resonance (DEER) spectroscopy has offered an excellent approach to obtain structural information on the conformational landscapes of complex protein systems that may be too disordered or flexible for cryoEM or crystallization (20, 21). Together with site-directed spin-labelling (SDSL), this technique uses radical-containing probes, such as nitroxides, that selectively label cysteine residues (22, 23). DEER spectroscopy is a pulsed electron spin resonance (ESR) technique that provides accurate distances between electron spins in the range of 1.5-8 nm (24, 25). This technique has previously been used to study the visual transduction pathway and has provided insights on rhodopsin activation of transducin as well as how arrestin engages rhodopsin (26, 27, 28). DEER can provide insight into the conformational changes that occur upon the binding of interaction partners, such as those between the PDEγ subunits and Gα_T_. Furthermore, because DEER reports on the full ensemble of states, it enables characterization of the multiple conformations present in a dynamic system, allowing us to investigate the alternating motions indicated by PCA (9). By nitroxide labeling of the single wild-type cysteine (PDEγC68-SL) on PDEγ as well as a cysteine residue introduced into a PDEγ (Ile64Cys, Cys68Ser) double mutant (PDEγC64-SL), we monitored the distances between the two PDEγ subunits under various conditions of Gα_T_* binding. DEER studies, as well as SAXS experiments, show that the binding of Gα_T_* to PDE6 causes extensive changes to the PDEγ subunits compared to the apo-state and that these changes differ dramatically when the active site is occupied by a slow hydrolyzing analog of the native substrate cGMP (8-Br-cGMP) compared to a substrate-competitive inhibitor. Importantly, the PDEγ-subunits appear much more loosely associated with the enzyme core in the complex containing the substrate, suggesting that high mobility of the Gα_T_:PDE6γ moieties is required for PDE6 activation, in keeping with the proposed alternating-site mechanism.

## Results

### Generation of spin-labelled PDE6 and predicted distance measurements

To generate spin-labelled PDE6 for detecting conformational changes of the PDEγ subunits, spin labels were introduced into two forms of recombinant PDEγ, wild-type (WT) PDEγ, which was labeled at C68 (PDEγC68-SL), and a PDEγ (Ile64Cys, Cys68Ser) double mutant labeled at C64 (PDEγC64-SL). These cysteine residues were labeled by incubating recombinant PDEγ with MTSL (a highly reactive, thiol-specific spin-label containing a nitroxide radical) (29). Native PDE6 purified from bovine retina (SI Appendix, Fig. 1A) was treated with trypsin, which selectively removes the native PDEγ subunits to generate a fully active PDE6 enzyme (tPDE6) (13). Trypsin-treated PDE6 (tPDE6) was then mixed with 2-equivalents of the spin-labeled PDEγ subunits to generate either PDE6:PDEγC68-SL or PDE6:PDEγC64-SL, the symmetrically labelled PDE6 holoenzymes (SI Appendix, Fig. 1B). These symmetrically labelled PDE6 holoenzymes were then assayed for their ability to be activated by the constitutively active recombinant Gα_T_* relative to the native WT PDE6. The basal activities of both spin-labeled holoenzymes in the absence of Gα_T_* are identical to that of the native enzyme, indicating that the spin-labelled PDEγ are fully functional and able to bind and inhibit the PDEα/PDEβ heterodimer. PDE6:PDEγC68-SL and PDE6:PDEγC64-SL stimulated nearly identical levels of cGMP hydrolysis at increasing concentrations of Gα_T_* and gave rise to slightly higher rates of catalytic activity compared to control enzyme (SI Appendix, Fig.1C). This likely reflects a slight decrease in PDEγ association with the PDEα/PDEβ heterodimer due to the flexibility of the spin-label. Given that the spin-labeled holoenzymes are strongly activated by Gα_T_*, we sought to examine the distances between the spin-labels on the two PDEγ subunits under varying conditions of Gα_T_* binding.

**Fig. 1:**
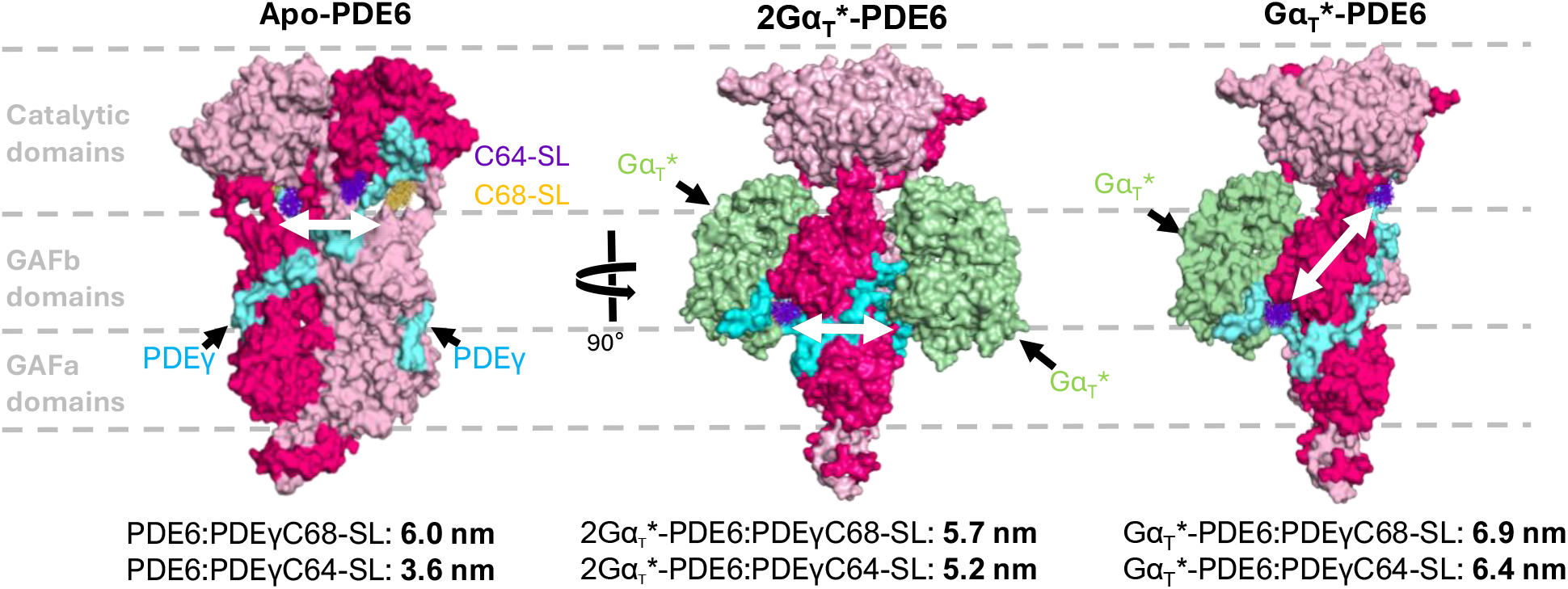
**Spin-label positions for structurally characterized PDE**γ **conformations.** Representative cryoEM models for each conformational state of the PDEγ subunits used for prediction of distance measurements by MMM (Multiscale Modeling of Macromolecules) (30). The predicted distances when labelled at Cys68 (PDE6:PDEγC68-SL) and Cys64 (PDE6:PDEγC64-SL) are shown for each conformation (PDE6 α-pink; β - dark pink; γ – cyan). The possible rotamers of the MTSL spin labels on C64 and C68 are shown in purple and yellow, respectively. Note that PDEγ is labelled at only either C68 or C64 for a given experiment. The constitutively active Gα_T_* subunits, which bind and rearrange the PDEγ subunits, are shown in green.

Prior to performing any DEER experiments, we utilized MMM (Multiscale Modeling of Macromolecules) (30) as a modeling tool to predict distances between spin labels on known cryoEM structures. For a given structural model and specified residue for labeling, this program evaluates all the possible rotamers of a given spin label. The populations of each rotamer are then computed, and the spatial distribution of the electron spins are modeled to predict the distance distributions between the two spins. These simulated distance distributions are shown in SI Appendix, Fig. S2. The structure of apo-PDE6 (PDB: 8ulg) (16) was predicted to have a distance of 6.0 ± 0.4 nm between spin labels placed on C68 (PDE6:PDEγC68-SL) (Fig. 1). The structure of the complex between PDE6 and two Gα_T_* subunits, which we refer to as 2Gα_T_*-PDE6:PDEγC68-SL (PDB: 7jsn) (9), predicted a distance of 5.7 ± 0.1 nm, with a much narrower distribution than the apo-state due to increased restraints on the spin-label upon the binding of GTP-bound Gα_T_* (Fig. 1). Additionally, an intermediate model wherein only one PDEγ subunit has been displaced from the catalytic domains by a single Gα_T_* (Gα_T_*-PDE6:PDEγC68-SL) was generated in PyMOL (31). This model predicted a longer spin-spin separation of 6.9 ± 0.4 nm (Fig. 1).

In 4-pulse DEER, it is important to note that reliably measured distances depend on t_max_ ≈ 1/(2 × dipolar coupling frequency), which in turn depends on the phase memory time of the spins (T_m_). Accurate distances at 6 nm separation (a dipolar coupling of ∼0.24 MHz) requires t_max_ of ∼4 μs or greater, whereas > 6 μs is required for the > 8 nm range (25). DEER data were collected with t_max_ = 6.5 μs, in keeping with potentially long spin-spin separation. Spin labels placed at residue 64 were predicted to produce distances much shorter than 6 nm and thus, PDE6:PDEγC64-SL was generated to gain additional information within the reliable distance range. The predicted separation distances for apo-PDE6:PDEγC64-SL, the 2Gα_T_*-PDE6:PDEγC64-SL complex, and the intermediate Gα_T_*-PDE6:PDEγC64-SL model, were 3.6 ± 0.3 nm, 5.2 ± 0.5 nm, and 6.4 ± 0.3 nm, respectively (Fig. 1, SI Appendix, Fig. S2B).

### Comparison of apo-PDE6 and PDE6 complexed to G*α*_T_*

Previous studies of the interaction between PDE6 and constitutively active Gα_T_* indicate that the binding of Gα_T_* causes extensive changes in the positions of the PDEγ subunits relative to the PDEα/PDEβ catalytic sites. To investigate these changes, we performed DEER experiments on PDE6:PDEγC68-SL and PDE6:PDEγC64-SL in the presence and absence of the constitutively active recombinant Gα_T_* subunit described previously (9). The spin labels on PDE6:PDEγC68-SL revealed a long, broad distance distribution with a mean value of 6.4 nm and a full width at half maximum (FWHM) of 2.3 nm (Fig. 2A). Interestingly, the distance was both longer and broader than what was predicted from the cryoEM structure for apo-PDE6 (PDB: 8ulg) (6.0 ± 0.4 nm). The distribution is skewed to larger separations with the predominant distances centering around 6.4 nm and trailing to > 9.0 nm; this long tail indicates that one or both of the spin labels are positioned further away from the PDEα/PDEβ catalytic sites than suggested by the cryoEM model. However, for experiments with a maximum evolution time of 6.5 μs, the shape of P(r) plots at distances greater than 7.0 nm are uncertain and their exact values should be interpreted with caution. PDE6:PDEγC64-SL gave a similarly broad DEER distance distribution with a mean of 5.1 ± 1.2 nm (FWHM) (Fig. 2A), much longer than the MMM predicted distance of 3.6 ± 0.3 nm. In this case, the distribution is skewed to shorter separations, with few distances close to the predicted distance of 3.6 nm and the largest population of inter-spin separations centered closer to 5-6 nm. Given that PDE6:PDEγC64-SL generates distance distributions within the reliable range of DEER, the shape of this P(r) plot can be reliably interpreted and indicates that one or both of the PDEγ subunits are flexible with a range of positions moving further away from the PDEα/PDEβ heterodimer than indicated by the cryoEM structure of apo-PDE6. When taken together with the fact that the PDEγ subunits are not fully resolved in most cryoEM structures of PDE6, these data suggest that the C-terminal ends of PDEγ have a high degree of flexibility and likely undergo rapid association-dissociation with the PDEα/PDEβ heterodimer in the apo-state. This apparent flexibility is supported by previous crosslinking studies indicating that PDEγ in the 64-68 region lacks secondary structure (8). Gα_T_ and the catalytic domains of PDEα/PDEβ bind to the same region of PDEγ (residues 63-87), however Gα_T_ has a stronger affinity (32, 33). The flexibility of PDEγ, together with the binding of ligands to the active sites, would enable Gα_T_ to productively bind and effectively out compete the catalytic domains of PDEα/PDEβ for the PDEγ subunits during enzyme activation, aiding in the rapid response associated with this signaling pathway. Furthermore, PDE6 has two distinct binding sites for PDEγ, each with different binding affinities (12, 8). Therefore, the broad distribution may arise from a state in which one PDEγ subunit binds to the higher affinity site and the second PDEγ is only transiently bound and flexible. The preclusion of two simultaneous tight binding interactions of PDEγ with the PDEα/PDEβ catalytic sites would be consistent with the proposed alternating-site mechanism (9).

**Fig. 2:**
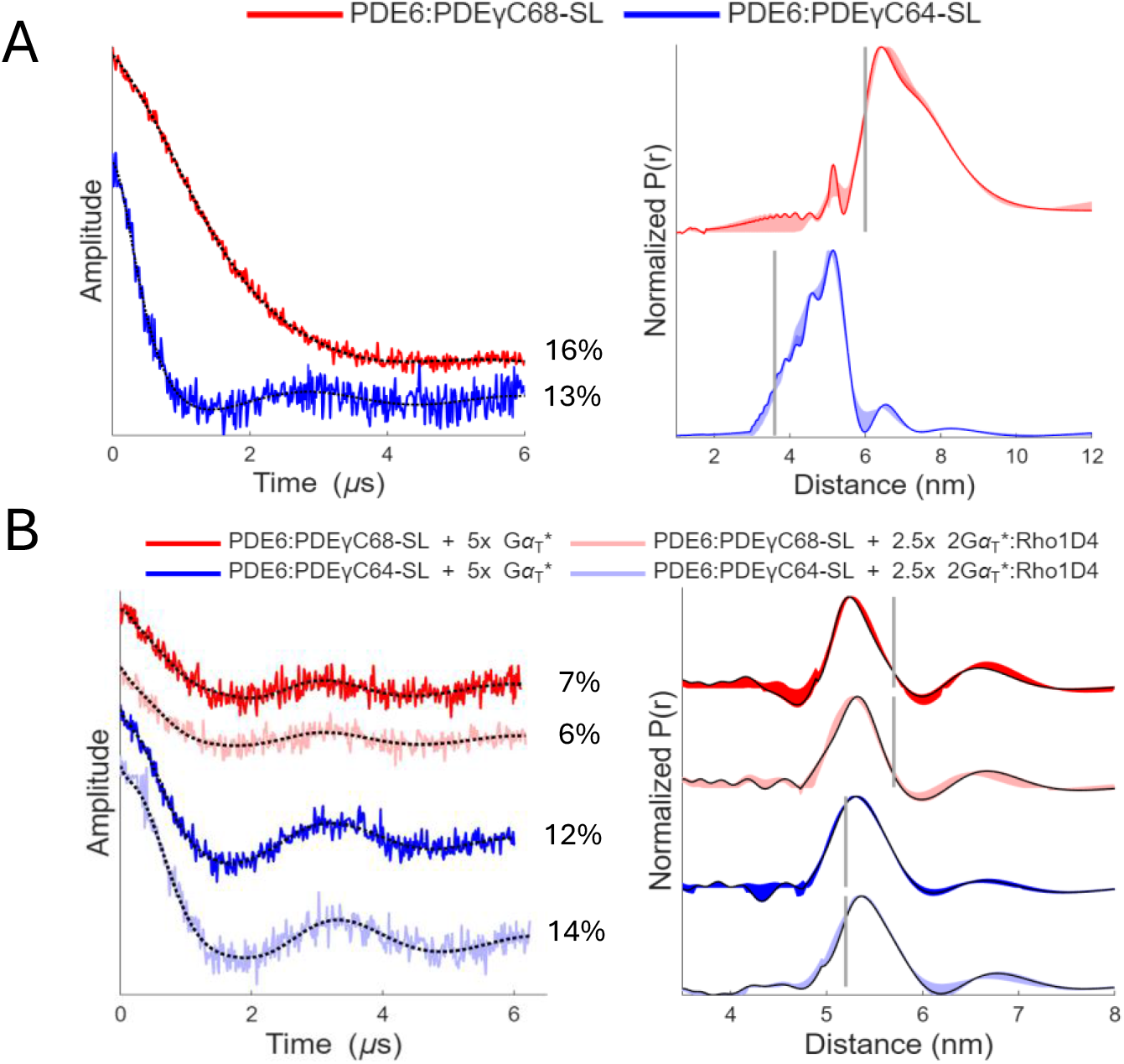
DEER spectroscopy of spin-labelled PDE6 in the presence and absence of. **G**α**_T_* and the bivalent antibody.** (A) Background subtracted DEER time domain traces (left) and corresponding experimental distance distribution plots (right) for PDE6:PDEγC68-SL (red) PDE6:PDEγC64-SL (blue). The MMM-simulated mean distances predicted for each labelling position on apo-PDE6 (PDB: 8ulg) are shown by the vertical gray line. Modulation depth percentage for each time domain trace is indicated to the right of each corresponding trace. (B) Background subtracted DEER time domain traces (left) and corresponding experimental distance distribution plots (right) for PDE6:PDEγC68-SL (red) and PDE6:PDEγC64-SL (blue) treated with udenafil and 5-fold excess Gα_T_* (5x monomer:PDE6 heterodimer) without the Rho1D4 bivalent antibody (dark) and with the 2Gα_T_*:Rho1D4 complex (light). The MMM-simulated mean distances predicted for each labelling position on the 2Gα_T_*-PDE6 model (PDB: 7jsn) are shown by the gray line. Modulation depth percentage is indicated to the right of each corresponding time domain trace.

Upon the addition of five-fold excess Gα_T_* (monomer-to-PDE6 heterodimer), together with the PDE6 inhibitor udenafil to occupy the active sites which is required for Gα_T_ binding (9, 16), DEER measurements of PDE6:PDEγC68-SL generated a distance distribution with the most prominent peak corresponding to a mean spin label separation of approximately 5.2 ± 0.5 nm (FWHM) (Fig. 2B). This distance agrees well with the MMM simulated distance between two spin-labeled PDEγ subunits at C68 on PDE6 (5.7 ± 0.1 nm), as determined from the 2Gα_T_*-PDE6:PDEγC68-SL cryoEM structure where both PDEγ subunits are displaced from the catalytic domains (Fig. 1). The slight discrepancy in the predicted and experimental distances is likely due to the favoring of spin-label rotamers that place the spins closer together than the average of all possible rotamers. Nevertheless, the experimental data agrees well with the cryoEM model. Another noteworthy feature is that despite preparing the samples identically, upon the addition of Gα_T_*, the DEER modulation depth produced by the sample decreased by about half (from 16% to 7%). Modulation depth represents the fraction of spins that are in the range for dipolar coupling and thus, about half of the spin-pairs in apo-PDE6:PDEγC68-SL are too far apart to be measured when in complex with Gα_T_*. This result could reflect an additional, broadly distributed state of the complex or partial dissociation of Gα_T_*:PDEγC68-SL. A second feature in the P(r) plot is enhancement of a peak centered at 6.5 nm, which is most prominent in all plots that demonstrate a decrease in modulation depth. This feature supports more flexible states in which Gα_T_* is interacting with PDEγ within the PDE6α/PDE6β heterodimer. To verify that the observed conformations were not altered by use of the constitutively active recombinant Gα_T_*, we measured udenafil-bound PDE6:PDEγC68-SL in the presence of excess native Gα_T_ loaded with slowly-hydrolyzing GTPγS. The resulting DEER trace is within error of that obtained with recombinant Gα_T_* (SI Appendix, Fig. S3). In the presence of udenafil and five-fold excess Gα_T_*, PDE6:PDEγC64-SL showed a mean inter-spin distance of 5.3 ± 0.5 nm (FWHM) which agrees well with the simulated distance of 5.2 ± 0.5 nm for the 2Gα_T_*-PDE6:PDEγC64-SL complex (Fig. 2B). We do not observe a decrease in modulation depth or the presence of a second peak in the P(r) plot of these samples.

Given that the only high-resolution structure of the interaction between PDE6 and Gα_T_* required the use of a bivalent antibody (Rho1D4) (34) that binds to the C-terminal end of Gα_T_* to stabilize the 2Gα_T_*-PDE6 complex, we examined whether the treatment of PDE6:PDEγC68-SL and PDE6:PDEγC64-SL with the antibody complexed to two Gα_T_* would produce a different result compared to the binding of Gα_T_* alone. Gα_T_* was incubated with the antibody to generate the 2Gα_T_*:Rho1D4 prior to mixing with spin-labelled PDE6 with Gα_T_* in five-fold excess (2.5-fold excess 2Gα_T_*:Rho1D4 complex). Treatment of both PDE6:PDEγC68-SL and PDE6:PDEγC64-SL with 2Gα_T_*:Rho1D4 and udenafil resulted in time domain traces and P(r) plots nearly identical to those with free Gα_T_*, 5.3 ± 0.5 nm and 5.4 ± 0.5 nm, respectively (Fig. 2B). These findings support the conclusion that the structure of 2Gα_T_*-PDE6 bound to udenafil was not significantly impacted by the use of this bivalent antibody (9).

### Titrations of G***α***_T_* with PDE6

We next sought to investigate the nature of Gα_T_* binding to PDE6 by monitoring the DEER signals with udenafil and different equivalents of Gα_T_* to spin-labelled PDE6. Each PDE6 hetero-tetramer has two Gα_T_ binding sites, one per each PDEγ subunit, and binding by Gα_T_ subunits at both sites is required for PDE6 activation (18). At 0.5x equivalents of Gα_T_* monomer to PDE6 heterodimer (a 0.25:1.0 subunit/subunit ratio), a PDE6:PDEγC68-SL spin-separation peak began to appear at 5.5 nm (Fig. 3A); at this ratio, a substantial population of distances representing apo-PDE6:PDEγC68-SL were present. At equal concentrations of Gα_T_* and spin-labelled PDE6 (1 Gα_T_* monomer to 1 PDE6 heterodimer), the distribution shifted toward a prominent distance with a mean at 5.3 nm, a similar distance to what was previously shown for PDE6:PDEγC68-SL treated with 5-fold excess Gα_T_* (i.e., assuming a 2 Gα_T_*: 1 PDE6 binding stoichiometry, Fig. 2B) with a population of distances > 6.0 nm still present. This second population of distances has a predominant curve ranging from 6.0 nm to 7.0 nm, as well as a shoulder indicating the presence of distances > 8.0 nm (Fig. 3A). We initially assumed that this second population corresponded to apo-PDE6; however, when a sufficient amount of Gα_T_* was added to occupy both binding sites (2 Gα_T_* monomer to 1 PDE6 heterodimer), the shoulder of distances > 8.0 nm disappeared, the modulation depth decreased by half, and a small peak with a mean of 6.5 nm remained present. This long distance peak and loss of modulation depth was still evident in our samples even in the presence of a 5-fold excess of Gα_T_* (5 Gα_T_* monomer to 1 PDE6 heterodimer) and could indicate the presence of a second population of dynamic states or the dissociation of some PDEγC68-SL from PDE6 upon Gα_T_* binding.

**Fig. 3:**
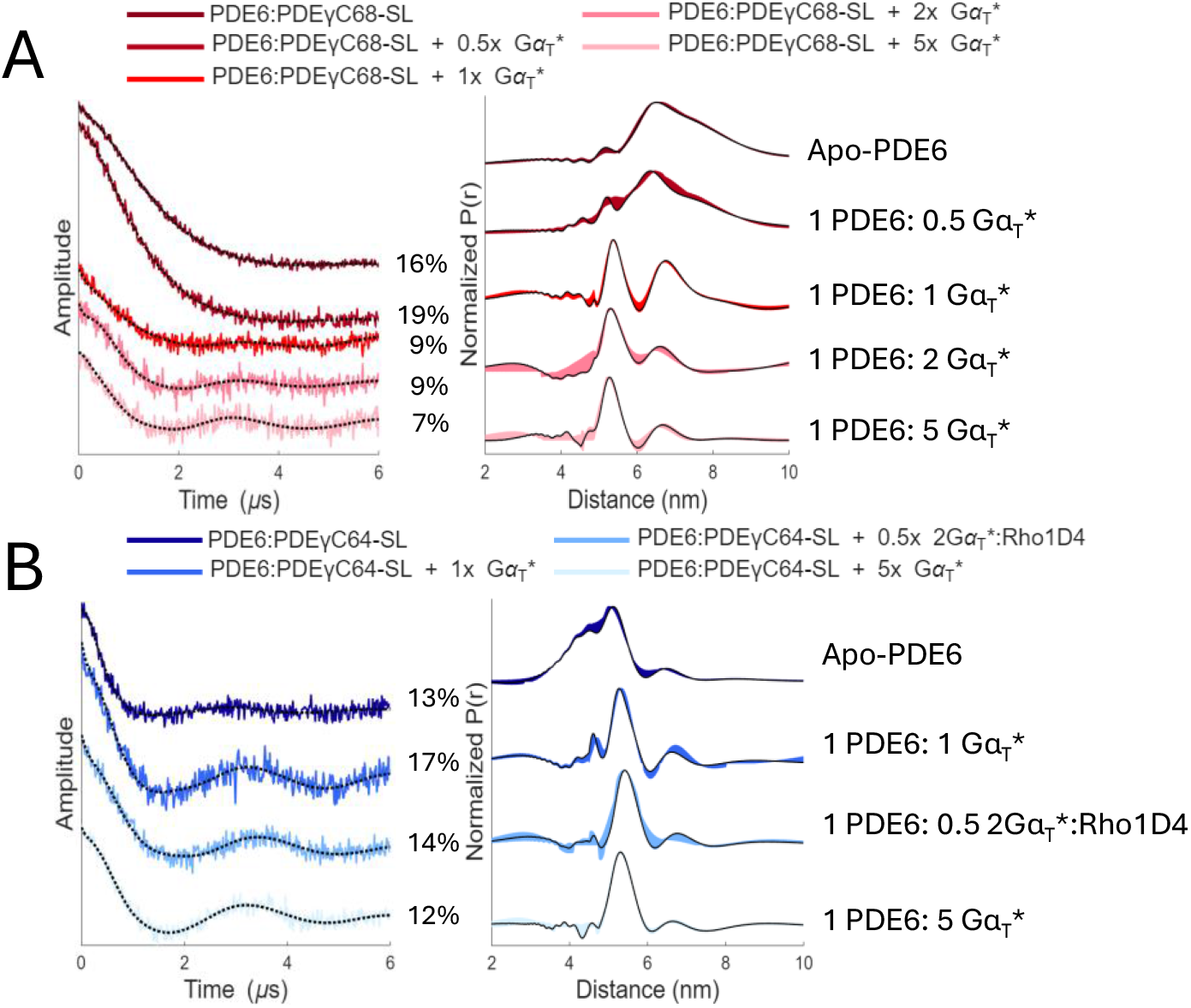
DEER-monitored titrations of spin-labelled PDE6 with increasing concentrations of. **G**α**_T_*.** Background subtracted DEER time domain traces (left) and corresponding distance distribution plots (right) for PDE6:PDEγC68-SL treated with increasing amounts of Gα_T_*. In descending order: apo-PDE6:PDEγC68-SL, 1 PDE6:PDEγC68-SL: 0.5 equivalents Gα_T_*, 1 PDE6:PDEγC68-SL: 1 equivalent Gα_T_*, 1 PDE6:PDEγC68-SL: 2 equivalents Gα_T_*, and 1 PDE6:PDEγC68-SL: 5 equivalents Gα_T_*. Modulation depth percentage is indicated to the right of each corresponding time domain trace. (B) Background subtracted DEER time domain traces (left) and corresponding distance distribution plots (right) for PDE6:PDEγC64-SL treated with sub-stoichiometric amounts of Gα_T_* with and without the bivalent antibody. In descending order: apo-PDE6:PDEγC64-SL, 1 PDE6:PDEγC64-SL: 1 equivalent Gα_T_*, 1 PDE6:PDEγC64-SL: 0.5 equivalents 2Gα_T_*:Rho1D4, and 1 PDE6:PDEγC64-SL: 5 equivalents Gα_T_*. Modulation depth percentage is indicated to the right of each corresponding time domain trace.

At a ratio of 1 Gα_T_* monomer to 1 PDE6 heterodimer (i.e., an amount of Gα_T_* sufficient to occupy half of the active sites of PDE6), and given a statistical distribution of Gα_T_* binding, we would expect the sample to assume a 1:2:1 ratio of apo-PDE6: Gα_T_*:PDE6: 2Gα_T_*:PDE6. However, we observed that the distribution largely resembles that of the state in which both PDEγ subunits have been repositioned by Gα_T_* in a 2Gα_T_*:PDE6 ratio. Thus, it is likely that the binding of the first Gα_T_* subunit significantly enhances the interaction of the second Gα_T_* with PDE6. This effect was observed again when PDE6:PDEγC64-SL was treated with sub-stoichiometric levels of Gα_T_* (1x Gα_T_*). This distribution is composed of distances predominantly centered at 5.3 nm, agreeing with the 2Gα_T_*-PDE6:PDEγC64-SL model (Fig. 3B). If there was a prominent state in which one Gα_T_* was bound to PDE6:PDEγC64-SL and displacing a single PDEγ, we would expect to see a population of distances centered near 6.4 nm as predicted by MMM for a Gα_T_*-PDE6:PDEγC64-SL model (Fig. 1). However, only a negligibly small peak greater than 6.0 nm can be seen in the distance distribution, and it is a considerably smaller population of distances compared to that of the 2Gα_T_*-PDE6:PDEγC64-SL state. Furthermore, in the distance distribution of PDE6:PDEγC64-SL treated with 1x Gα_T_* when Gα_T_* is complexed to the bivalent antibody (0.5x equivalents of 2Gα_T_*:Rho1D4), there was no difference in the DEER time domain traces compared to when treated with 1x free Gα_T_* (Fig. 3B). Because the antibody binds two Gα_T_* subunits and simultaneously presents a Gα_T_* subunit to each of the two binding sites on PDE6 (9), we expect it to produce a 2Gα_T_*:PDE6 state and thus mimic a cooperative binding interaction. Given that the traces of PDE6:PDEγC64-SL when treated with free Gα_T_* and Gα_T_* complexed to the bivalent antibody are nearly identical, we can assume that a complex in which two Gα_T_* are bound to PDE6 is more favorable than a state in which only one Gα_T_* is bound, supporting previous findings (9, 18, 35).

### Interaction between PDE6 and G***α***_T_* in the presence of 8-Br-cGMP

Structural studies of the Gα_T_-PDE6 signaling complex in the presence of the enzyme substrate, cGMP, have been challenging due to the fast catalytic turnover of cGMP and that upon the dissociation of the product GMP, the affinity of Gα_T_ for PDE6 is reduced (16). For this reason, we decided to utilize a slowly-hydrolyzing analog of cGMP, 8-bromo-guanosine 3′,5′-cyclic monophosphate (8-Br-cGMP) (18, 36), to determine if PDE6 induces conformational changes during substrate hydrolysis that are not detected in samples utilizing substrate competitive inhibitors for active site occupation. The maximal rate of 8-Br-cGMP hydrolysis is approximately 7 molecules per second, which is markedly slower than the maximal rate for cGMP hydrolysis, estimated at ∼4000 cGMP molecules hydrolyzed per second (37).

Addition of 25 mM 8-Br-cGMP to the spin-labelled enzyme alone produced a striking change in the time domain signals for both PDE6:PDEγC68-SL and PDE6:PDEγC64-SL compared to the apo-state (Fig. 4A, C). Given that apo-PDE6:PDEγC68-SL produces an already long distance even in the absence of 8-Br-cGMP, we also measured PDE6:PDEγC64-SL with 8-Br-cGMP. Upon addition of 8-Br-cGMP, background subtraction becomes difficult and less reliable as the spin separations become very long and distributed in both cases. Therefore, for samples treated with 8-Br-cGMP, analyzing the non-background corrected data (Fig. 4A, C) is helpful. Whereas a narrow distribution of distances leads to clear oscillations, in a broad range of distances, the frequencies will average out, producing a smooth monotonic decay (25). Comparing this signal to apo-PDE6:PDEγC64-SL, which has a prominent distance at 5.3 nm and clear oscillations in the signal, we can infer that the PDEγ subunits have moved far apart and are becoming less structured (Fig. 4C). This dramatic change in distance without the addition of Gα_T_* suggests that active site occupation by 8-Br-cGMP alone leads to a loss in the interaction of the C-terminal region of the PDEγ subunits with the PDE6 heterodimer. Given that active site occupation by substrate alone produces changes in the distance distribution between spin-labels, we also tested PDE6:PDEγC68-SL treated with udenafil. However, in contrast to the substrate, udenafil in the absence of Gα_T_* alters the time-domain trace in a manner consistent with a contraction in the spin label separation. Hence, unlike the substrate, the inhibitor stabilizes PDEγ against PDEα/PDEβ (SI Appendix, Fig. S4A). This stabilization effect leads to formation of a symmetric inhibited complex upon binding of Gα_T_* (SI Appendix, Fig. S4B).

**Fig. 4:**
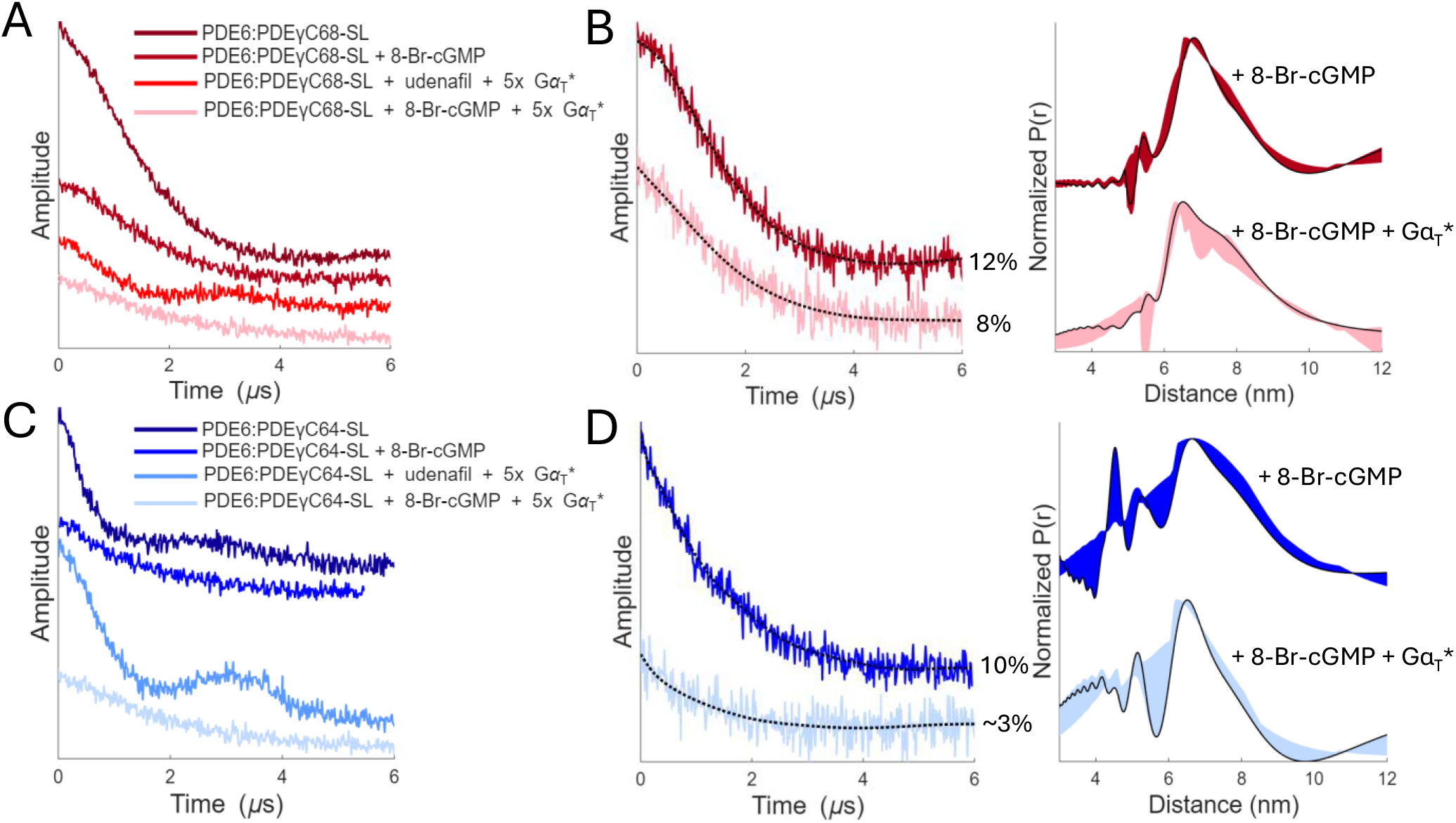
DEER spectroscopy of spin-labelled PDE6 treated with slowly hydrolyzing substrate 8-Br-cGMP and. **G**α**_T_*.** (A) Non-background-corrected time domain traces of PDE6:PDEγC68-SL under conditions of Gα_T_* binding with either udenafil or 8-Br-cGMP in the active site. In descending order, apo-PDE6:PDEγC68-SL, PDE6:PDEγC68-SL with 8-Br-cGMP, PDE6:PDEγC68-SL with udenafil and 5-fold excess Gα_T_*, and PDE6:PDEγC68-SL with 8-Br-cGMP and 5-fold excess Gα_T_*. (B) Background subtracted DEER time domain traces (left) and corresponding distance distribution plots (right) for PDE6:PDEγC68-SL with 8-Br-cGMP and PDE6:PDEγC68-SL with 8-Br-cGMP and 5-fold excess Gα_T_* (modulation depth percentage shown at right). (C) Non-background-corrected time domain traces of PDE6:PDEγC64-SL under conditions of Gα_T_* binding with either udenafil or 8-Br-cGMP in the active site. In descending order, apo-PDE6:PDEγC64-SL, PDE6:PDEγC64-SL with 8-Br-cGMP, PDE6:PDEγC64-SL with udenafil and 5-fold excess Gα_T_*, and PDE6:PDEγC64-SL with 8-Br-cGMP and 5-fold excess Gα_T_*. (D) Background subtracted DEER time domain traces (left) and corresponding distance distribution plots (right) for PDE6:PDEγC64-SL with 8-Br-cGMP and PDE6:PDEγC64-SL with 8-Br-cGMP and 5-fold excess Gα_T_* (modulation depth percentage shown at right).

Addition of both 25 mM 8-Br-cGMP and 5-fold excess Gα_T_* caused even larger increases in spin separation and distribution breadth compared to the substrate alone. For both PDE6:PDEγC68-SL and PDE6:PDEγC64-SL, the time domain traces were even more extended, less steep, and lost modulation depth, compared to both the apo-state and the inhibited complex (i.e., PDE6 bound to Gα_T_* with udenafil; Fig. 4A, C). PDE6:PDEγC68-SL treated with 8-Br-cGMP and excess retinal Gα_T_-GTPγS produces a similar DEER trace and distance distribution (SI Appendix, Fig. S3). With such distributed signals, the P(r) plots (Fig. 4B,D) indicate very heterogeneous conformational states. Given that at a maximum dipolar evolution time of 6.5 μs the maximum reliable distances are < 8.0 nm, distance separations beyond this limit are not accurate, but nevertheless indicate long and heterogenous distributions. This broad range of distances supports a highly dynamic state when both Gα_T_* and substrate is bound. Taken together, these experiments suggest that when the catalytic sites are occupied by 8-Br-cGMP, the Gα_T_* subunits cause rearrangements of the PDEγ subunits into a previously unobserved conformational state, which is much more flexible than when PDE6 is inhibited by the larger competitive inhibitors such as udenafil and vardenafil (9, 16).

### Small angle X-ray scattering

Given the highly dynamic state of PDE6 indicated by the DEER data in the presence of 8-Br-cGMP compared to udenafil, we investigated this structural change using small angle x-ray scattering (SAXS). Similar to DEER, SAXS is a technique that yields direct structural information on an entire conformational ensemble in solution and can report on conformational differences in flexible domains, such as the ones seen in PDEγ (38). Given the transient nature of Gα_T_ binding, we have been unsuccessful in isolating the 2Gα_T_*-PDE6 complex by size exclusion chromatography and therefore opted to use batch-mode SAXS to monitor how the overall sample composition changes when PDE6 and Gα_T_* are treated with excess 8-Br-cGMP or udenafil (Fig. 5). The scattering of PDE6 and Gα_T_* as individual components were first measured independently to ensure the samples were well behaved and not aggregated prior to mixing to form the 2Gα_T_*-PDE6 complex (SI Appendix, Fig. S5). Mixtures of PDE6 with 2-fold excess Gα_T_* were then measured in the presence of either udenafil or 8-Br-cGMP (SI Appendix, Fig. S6). The scattering profiles as *I(q) vs q* and the dimensionless Kratky representation (*qR_g_^2^ * I(q)/I*(*0*) *vs qR_g_*), which utilizes the mid-q region of the SAXS scattering and provides information on the shape, compactness, and flexibility of a protein (39), are shown for solutions of PDE6 and 2-fold excess Gα_T_* in the presence of excess udenafil or 8-Br-cGMP (Fig. 5). The structural parameters are reported in SI Appendix, Table S1. For a sample of PDE6 and a 2-fold excess Gα_T_* with no ligand, PDE6 and Gα_T_* are assumed to not interact and thus the scattering represents a combination of the scattering profiles of the two proteins individually. Upon addition of substrate or inhibitor, a majority of the free Gα_T_* binds PDE6, leading to changes in the mid-q region of the scattering that are apparent in the Kratky representation. All three curves have a prominent peak with a maximum at qR_g_ = ∼2, slightly shifted from that expected for a globular protein (√3, 1.104), and a characteristic shoulder. For the curve representing the scattering of free PDE6 and Gα_T_* in solution (PDE6 + 2x Gα_T_*), this shoulder is subtle and at a qR_g_ of ∼6.7, slowly decaying at higher q. When udenafil is added, the Kratky plot exhibits a narrower peak and a more defined shoulder at qR_g_ of ∼5.5, indicating a more compact and less flexible Gα_T_*-bound state with less conformational heterogeneity, consistent with DEER experiments in the presence of udenafil. In contrast, when 8-Br-cGMP is added, the scattering results in a broader curve compared to the udenafil-bound state that plateaus at high qR_g,_ consistent with an extended Gα_T_*-bound state with increased flexibility or partial disorder of the Gα_T_*:PDE6γ moieties (40). These data indicate that binding of 8-Br-cGMP promotes an extended and dynamically heterogeneous Gα_T_*-bound state. Furthermore, the simulated scattering curve of the 2Gα_T_*-PDE6 model computed by FoXS (41, 42) fits much closer to the udenafil-bound scattering curve (χ^2^ = 1.68) compared to that of the 8-Br-cGMP-bound state (χ^2^ = 5.27) (SI Appendix, Fig. S7), further supporting the idea that 8-Br-cGMP promotes a previously uncharacterized dynamic active state of the enzyme that is different from the cryoEM model.

**Fig. 5:**
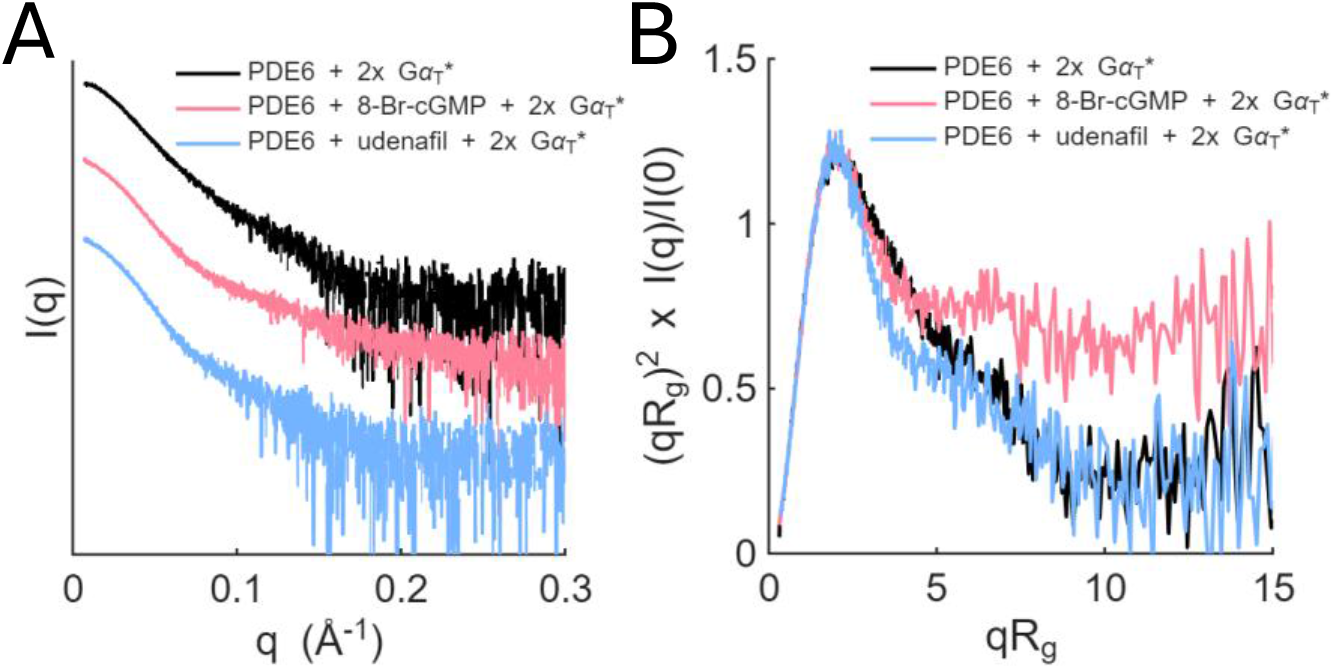
Small Angle X-ray scattering (SAXS) of Gα_T_* binding to PDE6 in the presence of 8-Br-cGMP or udenafil. Batch-mode SAXS data presented as (A) semilog plots and (B) overlayed dimensionless Kratky plots of PDE6 treated with 2-fold excess Gα_T_* either with the active sites unoccupied (black) or in the presence of 1 mM 8-Br-cGMP (pink) or 3-fold excess udenafil (blue). All three Kratky curves have a prominent peak with a maximum at qR_g_ = ∼2, slightly shifted from that expected for a globular protein (√3,1.104). Scattering curves on the semilog plots were offset for visual clarity and the dimensionless Kratky plot was re-binned to smooth noise. All structural parameters (R_g_, I(0)) were derived from unbinned data. The structural parameters and sample details can be found in SI Appendix, Table S1.

## Discussion

Upon activation by GPCRs, heterotrimeric G proteins dissociate into a GTP-bound Gα subunit and a Gβγ subunit complex that interact with effector enzymes to regulate intracellular second messengers such as cAMP and cGMP, thereby converting extracellular signals into amplified intracellular responses that control a range of biological processes including metabolism, sensory perception, and neurotransmission. In the visual transduction pathway, regulation of cGMP by PDE6 in retinal rod cells exemplifies a highly amplified G protein–effector interaction and has served as a model for understanding GPCR signaling, although the precise mechanism by which activated Gα_T_ regulates PDE6 via its PDEγ subunits remains unclear. Recent cryoEM studies provide the first high-resolution structure of PDE6 in complex with constitutively active Gα_T_* subunits, revealing that binding of two Gα_T_ subunits induces a pronounced displacement of the PDEγ C-termini away from the catalytic domains of the PDEα/PDEβ heterodimer (9). However, achieving this 2Gα_T_-PDE6 structure required stabilization of the complex using both the substrate-competitive inhibitor vardenafil and a bivalent antibody to maintain simultaneous binding of two Gα_T_* subunits. Subsequent studies performed without the antibody were unable to resolve Gα_T_ or the full PDEγ subunits at high resolution (16), highlighting the dynamic and flexible nature of these proteins. ESR techniques enabled us to directly probe the conformational landscape of the protein in the absence of the antibody and with a slowly hydrolyzing substrate. Here we report the first ESR spin-labeling study of a G protein interacting with its effector enzyme and provide new insights into the structure and dynamics of the PDE6–transducin complex under activating conditions.

Our experiments on apo-PDE6 spin labelled at residues 68 (PDE6:PDEγC68-SL) and 64 (PDE6:PDEγC64-SL) reveal that when the PDE6 active sites are unoccupied, the PDEγ subunits have a high degree of flexibility, with surprisingly long spin separations (Fig. 2A). In every high-resolution structure of PDE6, the PDEγ subunits are not fully resolved, especially residues 40-75, which have a high degree of flexibility. Cys 68 and Ile 64 were previously shown to interact with the PDE6 heterodimer in the PDEβ binding site but not in the PDEα subunit binding site in crosslinking studies (8). Furthermore, this crosslinking study showed that these flexible regions of PDEγ do not bind PDE6 in an extended linear conformation like the N-terminal GAF-binding region (residues 10-37) and the C-terminal region (residues 80-87) that binds the catalytic sites and instead assume a more flexible random coil (8, 43). The broad DEER distributions are consistent with two distinct PDEγ binding environments on the PDE6 heterodimer, with one site maintaining a high degree of flexibility, consistent with earlier biochemical studies suggesting unequal binding affinities at the two catalytic sites (12). Hence, the relatively static cryoEM model of apo-PDE6 (PDB: 8ulg) is not a complete representation of the conformational landscape of the protein. The preclusion of two simultaneous tight binding interactions of PDEγ with the PDEα/PDEβ catalytic dimer is also consistent with the proposed alternating-site mechanism for enzymatic activity (9).

In the presence of Gα_T_* and the competitive inhibitor udenafil, the DEER distance distributions shift to values that closely match those predicted from the cryoEM structure of the 2Gα_T_*-PDE6 complex, demonstrating that this restructuring of the PDEγ C-termini bound to Gα_T_* represents the majority of the sample (Fig. 2B). Importantly, the addition of Gα_T_* bound to the Rho1D4 bivalent antibody did not significantly alter the measured distances, indicating that the antibody primarily stabilizes the complex without introducing significant conformational changes to the structure. Further support for this overall conformation is seen in low resolution densities of this complex in the absence of the bivalent antibody in which density for two Gα_T_ subunits can be observed near the GAFb domains (16). However, the decrease in modulation depth observed in some samples of PDE6 treated with Gα_T_* indicates that either there may be another state present in which Gα_T_* moves the PDEγ subunits further away than what is represented by the cryoEM structure or that Gα_T_* destabilizes the binding of PDEγ to PDEα/PDEβ and causes some dissociation. In either case, the data suggest that a rigid model of Gα_T_* binding may not be representative of all relevant conformational states. Furthermore, DEER-monitored titrations of Gα_T_* revealed that sub-stoichiometric amounts of Gα_T_* (1 Gα_T_* monomer to 1 PDE6 heterodimer; an amount of Gα_T_* sufficient to occupy half of the active sites of PDE6) drive a substantial population of PDE6 into a state resembling the conformation in which both PDEγ subunits have been repositioned by Gα_T_* (Fig. 3). The fact that this population exceeds the fraction expected assuming a purely statistical distribution points toward cooperative binding of Gα_T_. However, a full analysis of cooperative binding proved difficult due to lack of a recombinant expression system for PDE6. Nevertheless, these observations are consistent with previous biochemical evidence that two Gα_T_ molecules are required for maximal activation and suggests that the binding of two Gα_T_ to PDE6 is preferred over a singly bound state (8, 18, 44).

The most striking observations emerged from experiments performed in the presence of the slowly hydrolyzing substrate analog 8-Br-cGMP. In contrast to the inhibitor-bound complexes, the presence of this substrate analog caused a dramatic increase in the apparent separation of the PDEγ spin labels and produced DEER signals indicative of extremely broad distance distributions that extend beyond the reliable detection range of 1.5-8 nm (Fig. 4). 8-Br-cGMP has previously been shown to be an active substrate, although it hydrolyzes at a much slower rate than cGMP. Therefore, this dynamic state likely represents a stabilized version of the active enzyme. Our results suggest that while the PDEγ subunits are quite flexible within the apo-PDE6 complex, the occupation of the catalytic sites by substrate that occurs even in the absence of Gα_T_* promotes a state in which the PDEγ subunits have weakened affinity for the PDEα/PDEβ heterodimer core, likely to aid in their recognition by Gα_T_. Addition of Gα_T_* under these conditions further increases the separation and breadth of the spin pair distributions, thereby revealing a distribution of enzyme conformations that is fundamentally different from the inhibitor-stabilized state observed by both DEER and cryoEM. Evidence for this dynamic active-state is further supported through the SAXS data demonstrating increased disorder and flexibility in samples with 2Gα_T_*-PDE6 treated with the substrate (Fig. 5). Taken together, our results suggest that Gα_T_ stimulates PDE6 catalytic activity not solely by displacement of the C-terminal ends of PDEγ from the catalytic domains as was thought previously (15), but instead likely through a multi-step alternating-site process (9). The first step involves substrate binding, which leads to a weakened interaction of PDEγ at the lower affinity PDEγ binding site with the catalytic core, thus allowing Gα_T_ to bind PDEγ and displace it to the GAF domains (Fig. 6). Binding of the first Gα_T_ subunit at the GAF domain then allows for binding of the second Gα_T_ subunit to the other PDEγ subunit. In the presence of an inhibitor, this leads to a symmetric “staging” conformation represented by the previously solved cryoEM structure (PDB: 7jsn) (9) that stabilizes the activating Gα_T_:PDEγ moieties against the PDEα/PDEβ core and largely holds the catalytic domains in the same positions as found for apo-PDE6 (SI Appendix, Fig. S4B). However, when substrate occupies the active site, the Gα_T_:PDEγ moieties transiently dissociate from the GAF domains, such that Gα_T_ activates the enzyme one active site at a time in an alternating fashion, as indicated by the long and broad DEER signals with the slowly hydrolyzing substrate (Fig. 4). Although the DEER and SAXS data provide evidence for such a dynamic active state, they do not, on their own, define this state as being asymmetric. This functional asymmetry has been suggested in previous studies due to the apparent difference of binding affinities between the two Gα_T_ binding sites (18, 44, 45) and the disproportionately greater impact of β-subunit mutations on rod cell stability and function (46). The enhanced dynamics, taken together with these previous studies and the finding that PDE6 in complex with Gα_T_ only achieves 50% of the activity observed in the absence of PDEγ, and that two Gα_T_ are required for activation, strongly implicate a model in which conformational flexibility of the PDEγ subunits enables sequential activation of the two catalytic domains (Fig. 6).

**Fig. 6:**
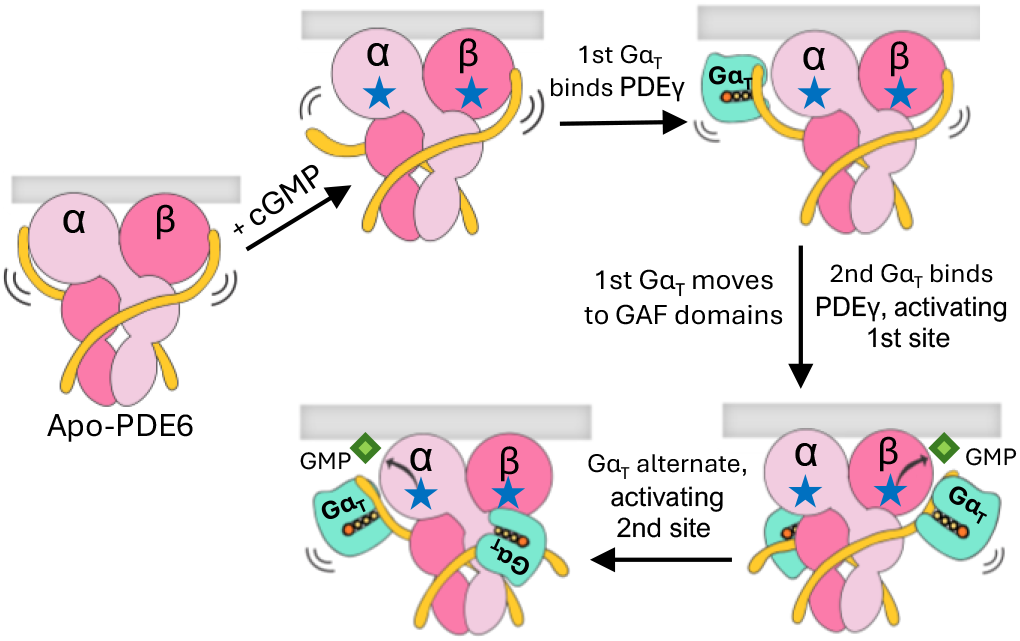
Conformational states associated with PDE6 activation. Flexible C-terminal ends of the PDEγ subunits (yellow) render the active sites of PDEα/PDEβ accessible to ligands. The binding of cGMP weakens the association of one PDEγ subunit with the catalytic domains of the PDEα/PDEβ core, increasing the distance between the PDEγ C-termini and aiding in the recruitment of Gα_T_. Upon rhodopsin activation of transducin, the GTP-bound Gα_T_ subunits rapidly associate with the flexible, dissociated PDEγ subunit and re-position the first PDEγ subunit to the GAF domains. Binding of this first Gα_T_ to the GAF domains relieves the inhibitory ability of the GAF domains and facilitates subsequent binding of the second Gα_T_ to the second PDEγ subunit. Upon binding of the second Gα_T_, cGMP hydrolysis occurs at the first active site. Sequential hydrolysis of cGMP at alternating catalytic sites is enabled by rapid transitions of Gα_T_ between GAF-associated and GAF-released states.

Our results indicate that PDE6 activation involves the restructuring of highly flexible PDEγ regulatory subunits that cannot be fully captured by static structural methods. Whereas inhibitors trap the 2Gα_T_*-PDE6 complex in a staging conformation with inactive catalytic domains, as represented by the cryoEM structure, the binding of the slowly hydrolyzing substrate 8-Br-cGMP favors a much more dynamic Gα_T_-promoted active state with loosely associated Gα_T_:PDEγ units, thereby relieving the ability of the GAF domains to inhibit catalysis. Interesting issues arise when this mechanism is considered in a biological context. For example, the cryoEM structure depicts each Gα_T_* subunit in the 2Gα_T_*-PDE6 complex oriented approximately 150° away from the assumed conformation of Gα_T_* when bound to rhodopsin on the membrane (9). Furthermore, PDE6 is prenylated on the C-terminal ends of its catalytic PDEα/PDEβ subunits and Gα_T_ is myristoylated on its N-terminal end (47). How these lipid modifications influence domain juxtaposition in the context of a membrane remain unknown, but ESR studies such as these provide an avenue to address such questions in more complex settings that involve membrane components.

## Conclusions

Overall, this study reveals new information regarding the mechanism by which GTP-bound Gα_T_ subunits activate PDE6 that were not revealed by our initial cryoEM structure of the Gα_T_-PDE6 complex. We demonstrate that the antibody does not influence the conformation of the inhibitor-bound Gα_T_-PDE6 complex. However, our new data shows that the symmetric doubly-bound Gα_T_-PDE6 state does not fully represent an active complex. We provide evidence that the tight regulation of cGMP hydrolysis required for vision in dim light occurs through a combination of steps that exceed the simple binding of Gα_T_ and movement of the PDE_γ_ subunits away from the active sites. This mechanism involves substrate binding to trigger the initial conformational changes that release the tight interaction of PDE_γ_ from the PDEα/PDEβ catalytic core, followed by the engagement of two Gα_T_ subunits with both PDE_γ_ and the GAF domains. The observed conformational heterogeneity indicates that these events occur in a coordinated yet asymmetric manner, consistent with an alternating-site mechanism in which the two catalytic centers are not simultaneously active. Whereas such dynamics had been previously hypothesized (9, 16), our results provide direct experimental support for their importance in catalysis and regulation. Indeed, the distinct differences between substrate-and inhibitor-bound states further suggest that this mobility is functionally coupled to catalysis, for example, with coordinated movement of Gα_T_ and PDE_γ_ enabling sequential turnover between active sites. Our working model (Fig. 6) takes into account previous findings and incorporates asymmetry and conformational dynamics as central features of this interaction.

## Methods

### Expression of recombinant 1D4-tagged Gα_T_* subunit

The plasmid encoding the recombinant Gα_T_* subunit containing an N-terminal 6xhis-tag and C-terminal 3-Ala linker connected to a Rho1D4 epitope tag (TETSQVAPA) and two GTP hydrolysis-defective mutations (R174C and Q200L) was generated as previously described (9). Recombinant Gα_T_* was transformed into BL21(DE3) competent cells and grown in six liters of TB media with 100 μg/ml ampicillin at 37 °C to an OD of 0.5. Protein expression was induced at 17 °C with 30 μM IPTG overnight.

### Purification of recombinant 1D4-tagged Gα_T_* subunit

The cells were resuspended in a lysis buffer (50 mM Tris pH 8, 500 mM NaCl, 5 mM MgCl2, 0.1 mM PMSF, 5 mM β-mercaptoethanol (BME), 10% glycerol) and lysed by sonication. The supernatant was collected after centrifugation at 185,000 x*g* for 45 minutes and then purified via a 5 mL HisTrap HP column (Cytiva Life Sciences) column equilibrated with 50 mM Tris pH 8, 20 mM imidazole, 5 mM BME. The loaded column was then washed with the equilibration buffer until the baseline was reached. The protein was then eluted with 50 mL of elution buffer (50 mM Tris pH 8, 20 mM imidazole, 5 mM BME, and 250 mM imidazole). The protein was further purified by anion-exchange chromatography using a 5 ml HiTrap Q HP column (Cytiva Life Sciences) equilibrated with buffer A (20 mM HEPES pH 7.5, 5 mM MgCl2, 10% glycerol) and eluted with a linear gradient to 50% buffer B (20 mM HEPES pH 7.5, 5 mM MgCl2, 10% glycerol, 1 M NaCl) in 150 mL. The eluted fractions were checked via SDS-PAGE, and the fractions containing the target protein were pooled and concentrated to about 50 μM, flash frozen, and stored at-80°C. Protein concentrations were determined using A_280_ and the extinction coefficient calculated with the ExPASy ProtParam tool to be 34,880 M^-1^cm^-1^.

### Purification of PDE6 from bovine retina

Retinal PDE6 was purified from bovine retina as previously described (16). Briefly, 300 dark-adapted bovine retina (InVision BioResources) were light-exposed, mechanically disrupted by shaking and homogenizing, and subjected to a series of centrifugations and sucrose gradient ultra-centrifugation steps to isolate the ROS disc membranes. The membranes underwent a series of wash steps, first with an isotonic buffer (10 mM HEPES pH 7.5, 100 mM NaCl, 1 mM DTT, 5 mM MgCl2, 0.1 mM EDTA), followed by three washes with hypotonic buffer (10 mM HEPES pH 7.5, 1 mM DTT, 0.1 mM EDTA) to extract peripherally associated PDE6 from the ROS membranes. PDE6 was then purified from the hypotonic wash by anion exchange chromatography on a 1 mL HiTrap Q HP column (Cytvia Life Sciences). The column was first equilibrated with buffer A (20 mM Tris pH 8.0, 5 mM MgCl2, 10% glycerol) and the protein was eluted by linear gradient to 50% buffer B (20 mM Tris pH 8.0, 5 mM MgCl2, 10% glycerol, 1 M NaCl). PDE6 was then concentrated to 10-15 μM with a 100 kDa MW cut-off centrifugal filter unit (Amicon, Millipore Sigma), flash frozen in liquid nitrogen, and stored at −80 °C.

### Cloning and mutagenesis of recombinant PDEγ

The DNA sequence encoding the 87 residue amino acid sequence of bovine retinal PDEγ (MNLEPPKAEIRSATRVMGGPVTPRKGPPKFKQRQTRQFKSKPPKKGVQGFGDDIPGMEGLGTDITVICPWEAFNHLELHELAQYGII (Uniprot ID: P04972)) was inserted into a pET-21(+) vector (Twist Biosciences) containing an ampicillin resistance marker. This DNA was used to generate the PDEγ Cys68Ser, Ile64Cys double mutant by first mutating the Cys68 to serine using the forward primer 5’-ATCACCGTTATCTCCCCATGGGAG-3’ and reverse primer 5’-CTCCCATGGGGAGATAACGGTGAT-3’. Mutagenesis of the DNA was then confirmed by sanger sequencing. Using the Cys68Ser mutant DNA, Ile64 was then mutated to a cysteine using the forward primer 5’-CTGGGTACCGACTGTACCGTTATCTCA-3’ and reverse primer 5’-TGAGATAACGGTACAGTCGGTACCCAG-3’. Mutagenesis of both point mutations were then confirmed by sanger sequencing.

### Expression and spin labelling of recombinant PDEγ

Wild-type PDEγ and the PDEγ Cys68Ser, Ile64Cys double mutant were expressed in BL21(DE3) cells by growing cells until an OD of 0.8-1 with 100 ug/mL ampicillin, followed by induction with 0.7 mM IPTG and continued cell growth for 4 hours at 37 °C. Harvested cells were then lysed by sonication in buffer A (25 mM Tris pH 7.5, 50 mM NaCl, 5 mM EDTA, 1 mM DTT) containing 1mM PMSF, 10 μM leupeptin, and 1 μM pepstatin A. The lysate was then clarified by ultracentrifugation at 185,000 xg for 45 minutes. The supernatant was injected onto a HiTrap SP HP cation exchange column (Cytiva Life Sciences) and eluted using a linear gradient to 100% buffer B (25 mM Tris pH 7.5, 1 M NaCl, 5 mM EDTA, 1mM DTT) in 150 mL. The peaks corresponding to PDEγ were isolated and concentrated to 5 mL using a 3 kDa molecular weight cut off Amicon Ultra-15 centrifugal filter device (Amicon, Millipore Sigma) and injected onto a HiLoad Superdex 75 pg size exclusion column equilibrated with 25 mM Tris pH 7.5, 150 mM NaCl, and 5% glycerol for further purification. The fractions containing purified PDEγ were then assessed for purity by SDS-PAGE, pooled, and concentrated to < 5 mL. Five-fold molar excess of (1-Oxyl-2,2,5,5-tetramethylpyrroline-3-methyl) methanethiosulfonate (MTSL) was added, the sample was diluted to 5 mL and then mixed overnight at 4°C. Unreacted MTSL was removed using a 5 mL Zeba desalting column (ThermoFisher). The desalted, spin-labelled protein was then assessed for concentration by A_280_ and the labelling efficiency was determined using continuous-wave ESR.

### Preparation of reconstituted PDE6 DEER samples

Five μM PDE6 in assay buffer (10 mM Tris pH 8.0, 2 mM MgCl2, and 100 mM NaCl) was trypsin-treated by the addition of 55 μg/mL TPCK-treated trypsin and incubation at room temperature for 3 minutes. At the end of the incubation period, 600 μg/mL soy-bean trypsin inhibitor (SBTI) was added to quench the reaction. The sample was then washed 3 times with assay buffer in a 0.5 mL 100 kDa molecular weight cut off centrifugal filter (Amicon, Millipore Sigma) to remove trypsin and SBTI and residual DTT from purification. A 3-fold molar excess spin-labelled PDEγ or the PDEγ Cys68Ser, Ile64Cys double mutant was added and allowed to incubate for 30 minutes on ice to reconstitute tPDE6 with spin-labelled PDEγ subunits and then washed with assay buffer to remove excess PDEγ. After reconstitution, 10-fold molar excess udenafil was added along with varying amounts of Gα_T_* or retinal Gα_T_ loaded with GTPγS. The samples were then buffer exchanged into 10 mM Tris pH 8, 100 mM NaCl, and 25% d8-glycerol buffer prepared in D_2_O using a 0.5 mL 10 kDa MW cut-off centrifugal filter unit, concentrated to ∼40 μM, and transferred to a 100 mm quartz Q-band EPR tube, checked via CW ESR, and flash frozen in liquid N_2_. Samples containing 8-Br-cGMP were prepared by first mixing spin-labelled PDE6 with 5-fold molar excess Gα_T_* or retinal Gα_T_-GTPγS, buffer exchanging and concentrating to ∼40 μM and then spiking with 25 mM 8-bromoguanosine 3′,5′-cyclic monophosphate (8-bromo-cGMP). The samples were quickly flash frozen in liquid N_2_ after the addition of substrate. For samples containing Gα_T_* bound to the Rho1D4 bivalent antibody (2Gα_T_*:Rho1D4 complex), Rho1D4 monoclonal antibody (University of British Columbia, CA, (34)) was first mixed with Gα_T_* as previously described (9) to generate the 2Gα_T_*:Rho1D4 complex and then added to udenafil-treated spin-labelled PDE6 in 2.5-fold excess, giving a final ratio of 5-fold excess Gα_T_* to spin-labelled PDE6.

### PDE6 activity assays

Hydrolysis of cGMP by PDE6 was measured as a function of pH change as previously described (16) using a micro pH electrode with a SevenDirect pH meter (Mettler Toledo). In 200 μL of assay buffer containing 10 mM Tris pH 8.0, 2 mM MgCl_2_, and 100 mM NaCl, varying amounts of Gα_T_* (0.25-2 μM) were incubated with 50 nM PDE6. Upon reaching a stable baseline, 800 nmole of cGMP was added and the decrease in pH (in mV) was recorded for 100s. The buffering capacity of the mixture was obtained by adding 800 nmol NaOH, and the cGMP hydrolysis rate (mol/sec) was determined from the ratio of the initial slope of the pH change in mV/sec and the buffering capacity of the assay buffer (mV/mol). The activity of the reconstituted, spin-labeled PDE6 was normalized to the activity of wild-type PDE6 treated with 2 μM Gα_T_*, which was considered to be the maximal activation of PDE6 under these conditions (100% activity).

### ESR spectroscopy measurements

CW ESR spectroscopy experiments were carried out at X band (∼9.4 GHz) on a Bruker E500 spectrometer equipped with a ER4123SHQE resonator. All experiments were performed at room temperature with a modulation amplitude of 2 G and 100 kHz modulation frequency. The acquisition parameters were fixed to 15 dB (6.325mW) microwave power and 60 dB receiver gain. All spectra were collected with 1024 points with a sweep width of 120 G for 4-8 scans.

### 4P-DEER

For four-pulse DEER, all samples were exchanged into buffer containing 25% d8-glycerol and then plunge frozen into liquid N_2_. Measurements were carried out at Q-band (∼34 GHz) on a Bruker Elexys E580 spectrometer equipped with a 10 W solid-state amplifier (150 W equivalent TWTA) and an arbitrary waveform generator. Experiments were performed at 50 K in an EN 5107D2 cavity with a cryogen-free insert/temperature controller. Four pulse DEER (π/2-τ1-π-τ1-πpump-τ2-π-τ2-echo) was carried out with 16-step phase cycling. The pump and probe pulses were separated by 50 MHz. The observer π/2 and π pulses, as well as the π pump pulse, had a length of 30 ns. Each DEER signal was acquired for 24-48 hours. The max dipolar evolution time (t_max_) was set to 6.5 μs with 410 points to balance the minimum trace length required to measure long distance contributions and maximize the signal to noise ratio.

### MMM simulations

For MMM distance predictions of PDE6:PDEγC68-SL, the cryoEM structures of apo-PDE6 (PDB: 8ulg), the 2Gα_T_*-PDE6 complex (PDB: 7jsn) and the Gα_T_*-PDE6 model generated in PyMOL (31) were loaded into MMM (30). The Gα_T_*-PDE6 model was generated in PyMOL by overlaying the 8ulg and 7jsn structures and then removing one PDEγ from each structure and one Gα_T_* from the 2Gα_T_*-PDE6 complex to generate a state with one PDEγ displaced by one Gα_T_* subunit. For PDE6:PDEγC64-SL, the Cys68 on the PDEγ subunits of these three models was first mutated to a serine residue and Ile64 was mutated to a cysteine residue using the mutagenesis tool in PyMOL. Upon uploading each model into the MMM interface, the residues of interest were selected (either Cys68 for PDE6:PDEγC68-SL or Cys64 for PDE6:PDEγC64-SL), the rotamers for MTSL at the labeled cysteines were computed at cryogenic temperature, attached, and then the simulated DEER distribution curves were generated along with the mean and standard deviation.

### ESR spectroscopy analysis

The 6.5 μs DEER traces were trimmed to remove end artifact distortion, then background subtracted. Distance distributions were obtained by the SF-SVD method (48) and the distance domain was normalized by fixing the maximum probability to 1. Data processing of the SF-SVD-based method was used through the SVDReconstruction software (https://denoising.cornell.edu). Modulation depth was determined after background subtraction using DeerAnalysis (49). Uncertainty estimates were determined using the method described by Srivastava and Freed, 2019 (50).

### SAXS sample preparation

PDE6 and Gα_T_* were first purified by size-exclusion chromatography in buffer containing 25 mM Tris pH 8.0, 100 mM NaCl, 2 mM MgCl_2_, and 2% glycerol using a Superdex 200 10/300 GL column. PDE6 was then concentrated to about 5 μM and Gα_T_* to about 30 μM. Samples of PDE6 at 0.3 mg/mL with 2-fold excess Gα_T_* containing either 3 μM udenafil or 1 mM 8-Br-cGMP were prepared with a final volume of 30 μL and then centrifuged at 14,000 x*g* for 10 min at 4°C before loading. Identical buffer matches for each sample were measured and subtracted from the scattering data. For samples containing 8-Br-cGMP or udenafil, equivalent amounts of each ligand were added to the respective buffer samples for proper subtraction. Sample information can be found in SI Appendix, Table S1.

### SAXS data collection and analysis

Batch-mode experiments on PDE6 and Gα_T_* were performed at the 7A1 station of the Cornell High Energy Synchrotron Source (CHESS) with sample oscillation in an in-vacuum X-ray sample cell using an x-ray energy of 11.3 keV and an attenuated flux of 2.01 x 10 11 ph/s at the sample. The x-ray beam was defined by an aperture of 250 µm by 250 µm. Each sample was continuously oscillated at room temperature while scattering was acquired over 30 seconds, with 1 second exposure per data frame, and the detector collected data over a q-range of ∼0.006-0.5 Å^-1^. All data frames were reduced, averaged, and buffer subtracted with BioXTAS RAW (51). GNOM was used to calculate the pair distance distribution function and D_max_ (52). Model fitting was done with FoXS (41, 42). All structural parameters are reported in SI Appendix, Table S1.

## Acknowledgments

These studies were supported by the National Institute of Health award EY034867 from the National Eye Institute. Electron spin resonance experiments were conducted at ACERT which is supported by the National Institute of General Medical Sciences of the National Institutes of Health under Award Number 1R24GM146107. Small angle X-ray scattering experiments were conducted at the Center for High-Energy X-ray Sciences (CHEXS), which is supported by the National Science Foundation (BIO, ENG and MPS Directorates) under award DMR-2342336, and the Macromolecular Diffraction at CHESS (MacCHESS) facility, which is supported by award P30-GM124166 from the National Institute of General Medical Sciences and the National Institutes of Health.

