## Supplementary material for "A highly dynamic active state for transducin-bound phosphodiesterase-6 in vertebrate phototransduction": SI Appendix

607-254-8634

**This PDF file includes:**

Figures S1-S7

Table S1

SI references

SUPPLEMENTARY FIGURES

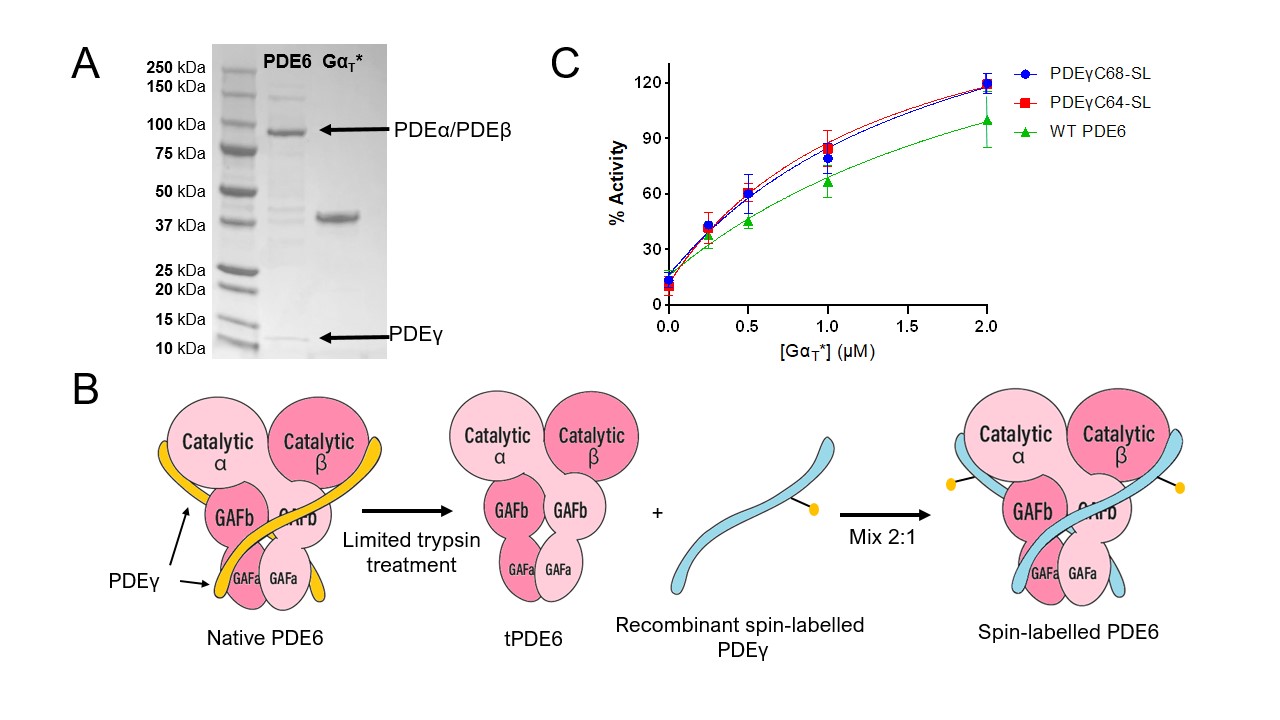

**Figure S1: Spin-labelling of PDE6 for ESR experiments.** (A) SDS-page gel of PDE6 purified from bovine retina prior to limited trypsin treatment (lane 1) and the recombinant Gα_T_* used for DEER experiments (lane 2). (B) Schematic depicting the reconstitution of PDE6 with spin-labelled PDEγ. Native PDEγ (yellow) is selectively digested with trypsin to generate tPDE6. tPDE6 is then mixed with a 2-fold excess of recombinant, spin labelled PDEγ (blue) to replace the native PDEγ and generate a symmetrically spin-labelled form of PDE6. (C) PDE6 activity assays comparing the relative activities of WT PDE6 and PDE6 reconstituted with PDEγ spin-labelled at the 64^th^ and 68^th^ positions. Percent activity is normalized to the average rate of cGMP hydrolysis of WT PDE6 in the presence of 2 µM Gα_T_*, which represents the maximum stimulated activity of PDE6 (100%). Experiments were performed in triplicate, and the standard deviation is represented as error bars.

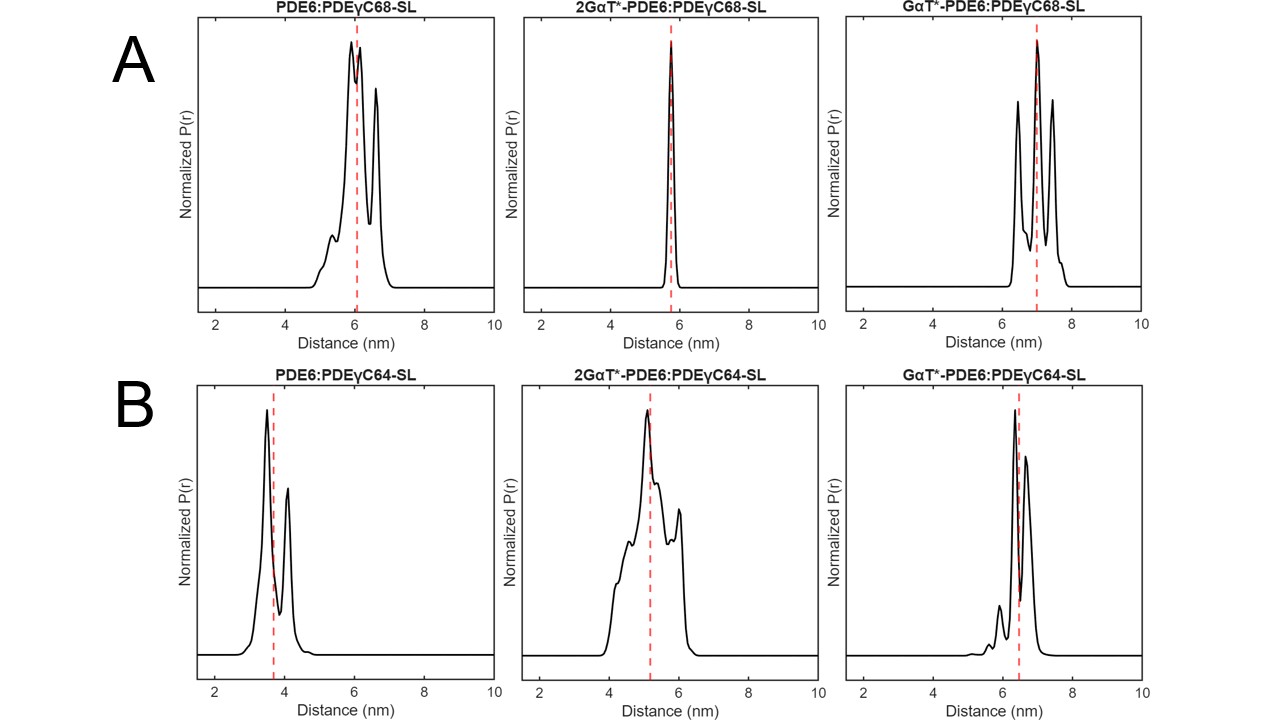

**Figure S2: Simulated distance distribution plots of the cryoEM structural models with Multiscale Modeling of Macromolecules (MMM)**. All simulations were carried out in MMM (1) by computing the possible rotamers for MTSL attached to the specified cysteine residues at cryogenic temperatures. The spatial distribution between all possible rotamers was then computed and plotted. (A) Simulated distributions for PDE6 spin labelled on Cys68 (PDE6:PDEγC68-SL) in the apo state (left), 2Gα_T_*-PDE6:PDEγC68-SL complex (middle) and Gα_T_*-PDE6:PDEγC68-SL model (right). (B) Simulated distributions for PDE6 spin labelled on Ile64Cys (PDE6:PDEγC64-SL) in the apo state (left), 2Gα_T_*-PDE6:PDEγC64-SL complex (middle) and Gα_T_*-PDE6:PDEγC64-SL model (right). For PDE6:PDEγC64-SL, Ile64 was first mutated to a cysteine residue in PyMOL (2) in all models prior to simulation.

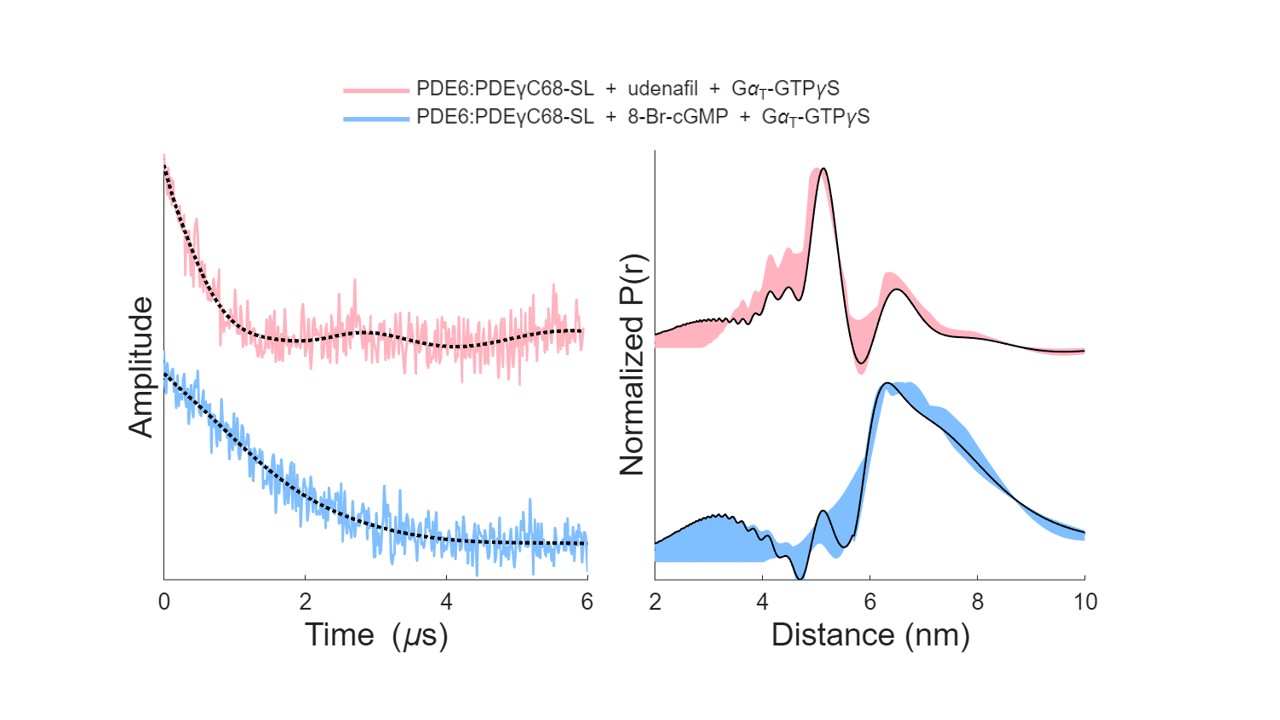

**Figure S3: DEER traces of PDE6:PDEγC68-SL treated with native retinal Gα_T_ bound to slowly hydrolyzing GTPγS in the presence of udenafil and 8-Br-cGMP.** Background subtracted DEER time domain traces (left) and distance distribution plots (right) of PDE6:PDEγC68-SL treated with natively purified retinal Gα_T_ loaded with GTPγS in the presence of udenafil (pink) and 8-Br-cGMP (blue). The P(r) plot of PDE6:PDEγC68-SL treated with Gα_T_-GTPγS and udenafil has a mean distance of 5.1 ± 0.5 nm, within error of that with recombinant Gα_T_* (5.2 ± 0.5 nm, Fig. 2B). The P(r) of PDE6:PDEγC68-SL treated with Gα_T_-GTPγS and 8-Br-cGMP has a mean distance of 6.3 ± 2.2 nm, compared to 6.5 ± 2.6 nm for that with recombinant Gα_T_* (Fig. 4D).

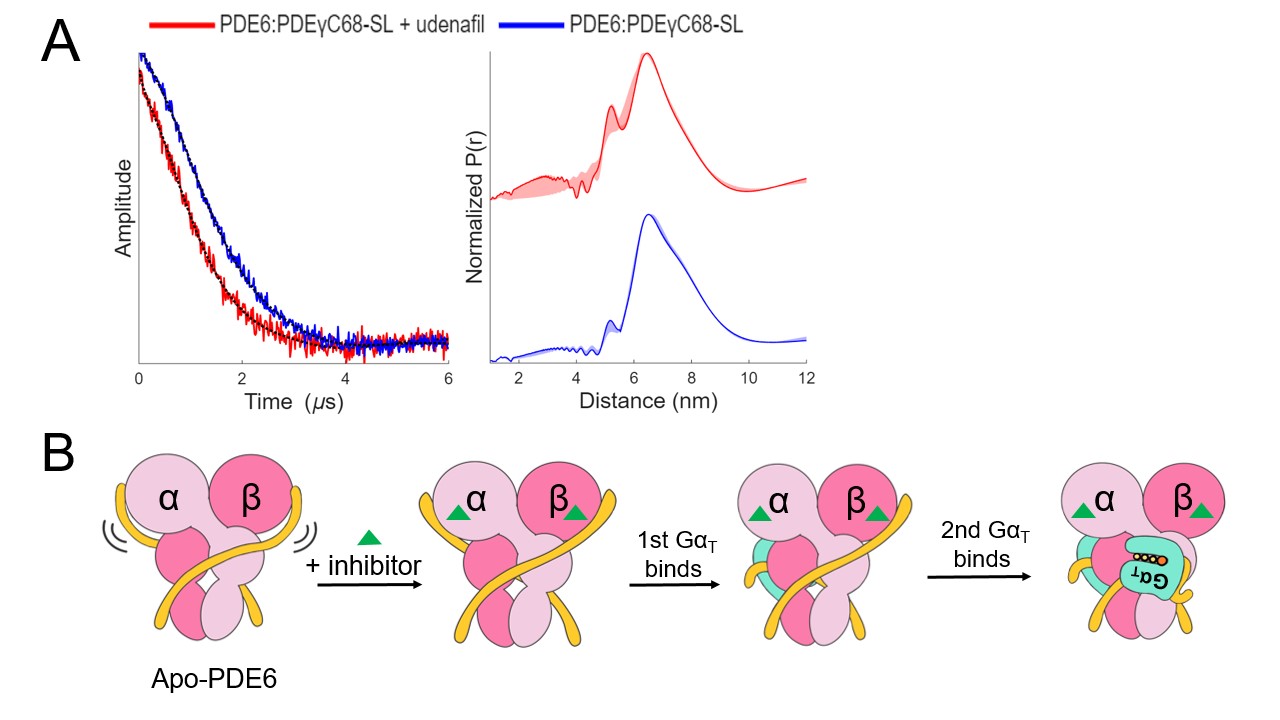

**Figure S4: The stabilizing effect of udenafil on spin-labelled PDE6.** (A) The background subtracted time domain traces (left) and distance distribution plots (right) comparing PDE6:PDEγC68-SL in the presence (red) and absence (blue) of udenafil indicate that the distances between spin labels decrease when udenafil occupies the active sites. When udenafil is present, the signal amplitude decreases more quickly, therefore indicating a shorter distribution of distances, and the P(r) plot correspondingly shows an increased prominence of a separations centered near 5.5 nm, indicating that the inhibitor has a stabilization effect on the PDEγ subunits. (B) Schematic demonstrating that the addition of large, competitive inhibitors such as udenafil stabilizes PDEγ on the enzyme but still allows binding of Gα_T_ and formation of a symmetric inhibited complex wherein the PDEγ subunits are pulled down beside the GAF domains, forming a state resembling the cryoEM model for the 2 Gα_T_*:PDE6 complex.

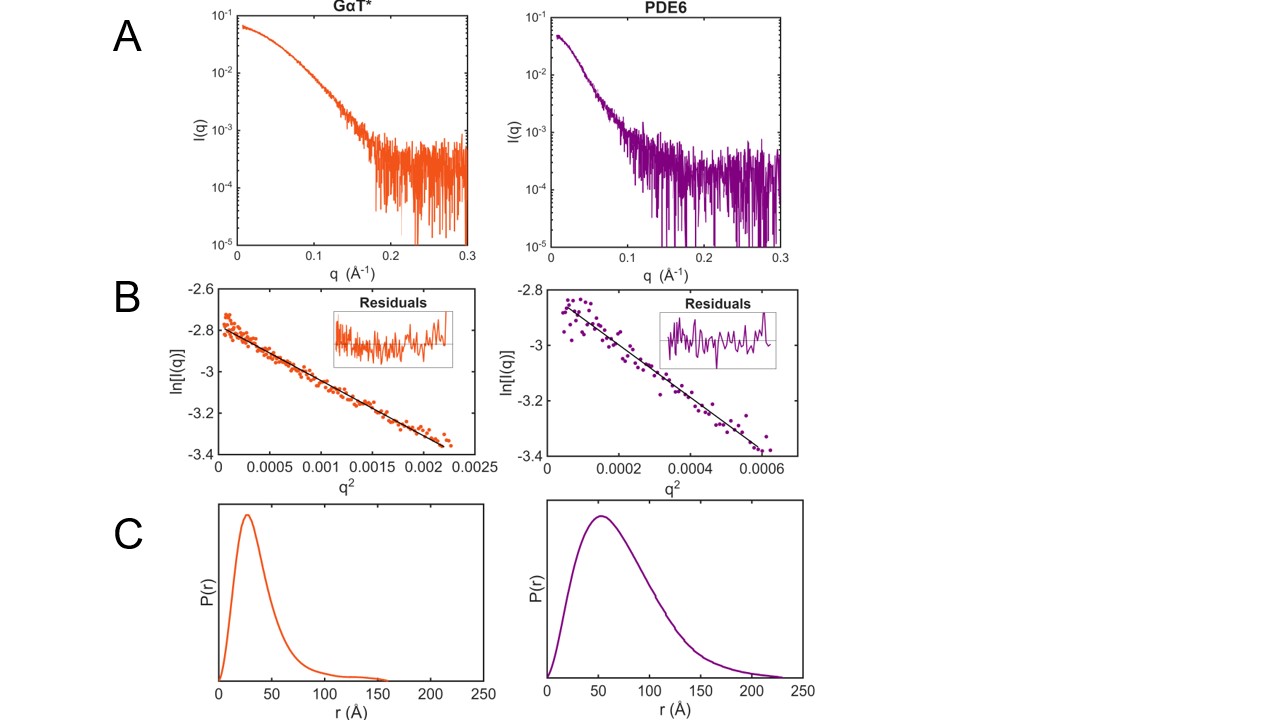

**Figure S5: Batch-mode SAXS data for Gα_T_* and PDE6 as individual components.** (A) SAXS data presented as semilog plots for Gα_T_* (orange) and PDE6 (purple). All data frames were reduced, averaged, and buffer subtracted with BioXTAS RAW (3). (B) Guinier fits for the SAXS data presented in panel A. Insets: normalized residuals for the respective fits. Both datasets have good Guinier regions, without deviation from linearity at low-q. (C) GNOM- calculated P(r) functions for the SAXS data (4). Data processing statistics can be found in Table S1. For both samples, up to 5 of the first data points were excluded from the scattering curve and Guinier fits to eliminate interference from the beamstop.

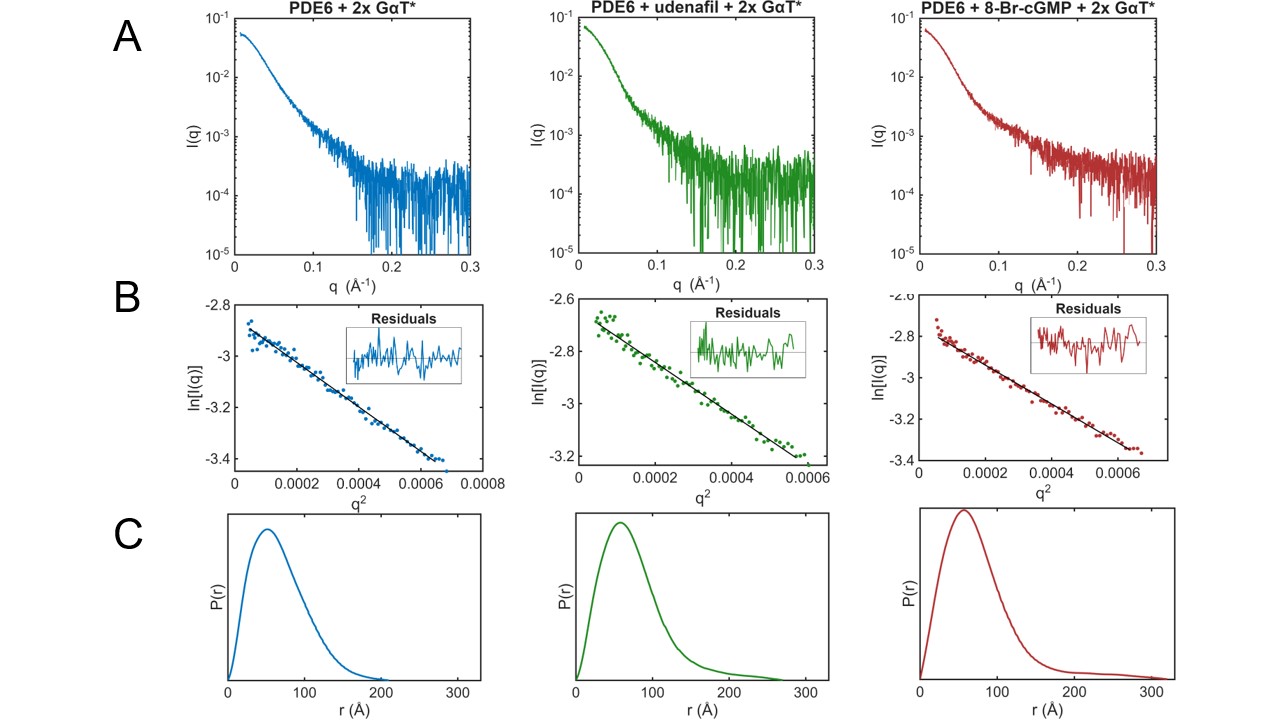

**Figure S6: Batch-mode SAXS data for the mixtures of PDE6 and Gα_T_* alone and in the presence of 8-Br-cGMP and udenafil.** (A) SAXS data presented as semilog plots for PDE6 and 2-fold excess Gα_T_* in the absence of ligand (blue), with 3 μM udenafil (green), and with 1 mM 8-Br-cGMP (red), respectively. All data frames were reduced, averaged, and buffer subtracted with BioXTAS RAW (3). (B) Guinier fits for the SAXS data presented in panel A. Insets: normalized residuals for the respective fits. All three datasets have expected Guinier regions, without deviation from linearity at low-q. (C) GNOM- calculated P(r) functions for the SAXS data (4). The shapes of the P(r) functions for all three datasets are highly similar; the slightly longer tails (and resulting higher D_max_ values) for the samples containing udenafil and 8-Br-cGMP coming from binding of Gα_T_*. The D_max_ of the sample containing 8-Br-cGMP is the largest due to the increased flexibility of the interaction when substrate occupies the active site. Data processing statistics can be found in Table S1. For all samples, up to 5 of the first data points were excluded from the scattering curve and Guinier fits to eliminate interference from the beamstop.

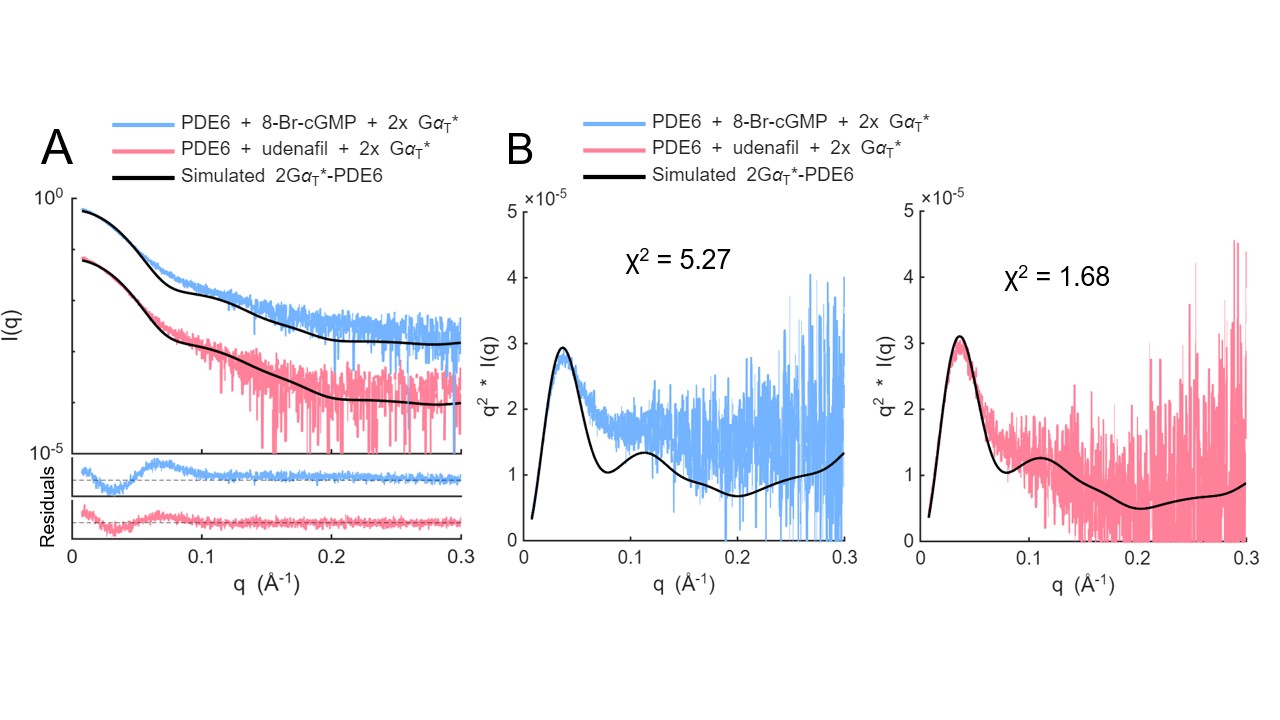

**Figure S7: Comparison of the 2Gα_T_*-PDE6 structural model with experimental scattering profiles.** (A) Experimental SAXS profiles of PDE6 with 2-fold excess Gα_T_* compared with the theoretical scattering profile (black) calculated from the 2Gα_T_*-PDE6 cryoEM structural model (PDB: 7jsn) using FoXS (5, 6). The residuals between experimental and simulated data are shown in the bottom panel and highlight differences in the mid-q region. There is much better agreement with the PDE6 + udenafil + 2x Gα_T_* scattering curve (χ^2^= 1.68) than the PDE6 + 8-Br-cGMP + 2x Gα_T_* scattering curve (χ^2^= 5.27), highlighting the differences in structure when substrate is present versus the inhibitor udenafil. (B) The scattering profile as a Kratky plot to highlight the structural differences between the two samples and the 2Gα_T_*-PDE6 structural model. The PDE6 + 8-Br-cGMP + 2x Gα_T_* Kratky plot highlights differences in the mid-q region, indicating a change in the domain positioning in the experimental data and flexibility not captured by the model. The PDE6 + udenafil + 2x Gα_T_* scattering curve has a smaller difference in the mid-q region when compared to the simulated curve, and it’s likely that this difference arises from heterogeneity of states in the sample solution rather than major structural differences in the 2Gα_T_*-PDE6 conformational state.

SUPPLEMENTARY TABLES

**Table S1**	SAXS sample details, data collection, and analysis.

| (a) Sample details |  |  |  |
| --- | --- | --- | --- |
| *Sample* | PDE6 (SASDZ96) | Gα_T_* (ID SASDZA6) |  |
| Source organism | *Bos taurus* | *Escherichia coli* |  |
| Protein | phosphodiesterase-6 | Constitutively active mutant of transducin, α-subunit |  |
| Buffer composition | 25 mM Tris pH 8.0, 100 mM NaCl, 2 mM MgCl2, 2% glycerol | 25 mM Tris pH 8.0, 100 mM NaCl, 2 mM MgCl2, 2% glycerol |  |
| Sample temperature (°C) | 4 | 4 |  |
| Sample concentration (mg/ml ) | 0.3 | 1.5 |  |
| (b) Sample details (mixtures) |  |  |  |
| *Sample* | PDE6 + 2x Gα_T_* (SASDZB6) | PDE6 + 2x Gα_T_* + udenafil (SASDZC6) | PDE6 + 2x Gα_T_* + 8-Br-cGMP (SASDZD6) |
| Buffer composition | 25 mM Tris pH 8.0, 100 mM NaCl, 2 mM MgCl2, 2% glycerol | 25 mM Tris pH 8.0, 100 mM NaCl, 2 mM MgCl2, 2% glycerol | 25 mM Tris pH 8.0, 100 mM NaCl, 2 mM MgCl2, 2% glycerol |
| Sample temperature (°C) | 4 | 4 | 4 |
| Sample concentrations (mg/ml) | 0.3 mg/mL PDE6 0.13 mg/mL Gα_T_* | 0.3 mg/mL PDE6 0.13 mg/mL Gα_T_* | 0.3 mg/mL PDE6 0.13 mg/mL Gα_T_* |
| Ligand |  | udenafil | 8-Bromoguanosine-3',5'-cyclic monophosphate |
| Ligand concentration |  | 3 μM | 1 mM |
| (c) Data collection |  |  |  |
| Beamline/Detector | CHESS 7A1 / Eiger 4M |  |  |
| Measured *q*-range (1/Å) | 0.006-0.5 |  |  |
| Detector distance (m) | 1.848 |  |  |
| Number, exposure time | 30 x 1 s |  |  |
| Energy (keV) | 11.3 |  |  |
| Flux at sample (ph/s) | 2.01 x 10^11 |  |  |
| Beam size (µm) | 250 x 250 |  |  |
| Experiment temperature (°C) | 4 |  |  |
| Configuration | batch |  |  |
| (d) Structural parameters |  |  |  |
| Sample | PDE6 | Gα_T_* |  |
| *Guinier Analysis* |  |  |  |
| I(0) ± s | 0.06 ± 0.0005 | 0.06 ± 0.0002 |  |
| R_g_ ± s (Å) | 53.3 ± 0.7 | 28.3 ± 0.2 |  |
| qR_g_ range | 0.4 -1.3 | 0.22 - 1.3 |  |
| Linear fit assessment (R2) | 0.955 | 0.982 |  |
| *GNOM/P(r) analysis* |  |  |  |
| I(0) ± s | 0.06 ± 0.0006 | 0.06 ± 0.0003 |  |
| R_g_ ± s (Å) | 57.9 ± 0.82 | 32.4 ± 0.6 |  |
| d_max_ (Å) | 230 | 160 |  |
| q-range (Å^-1^) | 0.0075 - 0.3 | 0.0076 - 0.3 |  |
| P(*r*) fit χ2/total estimate | 0.81778/0.7276 | 0.7928/0.6363 |  |
| MW (kDa) (Vc) | 218.1 | 45.1 |  |
| (e) Structural parameters contd. |  |  |  |
| Sample (mixtures) | PDE6 + 2x Gα_T_* | PDE6 + 2x Gα_T_* + udenafil | PDE6 + 2x Gα_T_* + 8-Br-cGMP |
| *Guinier Analysis* |  |  |  |
| I(0) ± s | 0.06 ± 0.0002 | 0.07 ± 0.0004 | 0.06 ± 0.0003 |
| R_g_ ± s (Å) | 51.08 ± 0.3 | 54.4 ± 0.5 | 53.1 ± 0.3 |
| qR_g_ range | 0.34 - 1.3 | 0.35 - 1.3 | 0.4 - 1.3 |
| Linear fit assessment | 0.9856 | 0.9784 | 0.9895 |
| *GNOM/P(r) analysis* |  |  |  |
| I(0) ± s | 0.06 ± 0.0003 | 0.07 ± 0.0006 | 0.07 ± 0.0006 |
| R_g_ ± s (Å) | 53.6 ± 0.43 | 60.4 ± 1.1 | 62.9 ± 1.5 |
| d_max_ (Å) | 210 | 270 | 320 |
| q-range (Å^-1^) | 0.007 - 0.3 | 0.007 - 0.3 | 0.008 - 0.3 |
| P(*r*) fit χ2/total estimate | 0.7356/0.7683 | 0.8217/0.6735 | 0.8389/0.6511 |
| (f) Model fitting |  |  |  |
| Experimental scattering curve | PDE6 + 2x Gα_T_* + udenafil | PDE6 + 2x Gα_T_* + 8-Br-cGMP |  |
| Program | FoXS | FoXS |  |
| Model | PDB: 7JSN | PDB: 7JSN |  |
| Chi-squared (χ2) | 1.68 | 5.27 |  |
| q-range (1/Å) | 0.006 - 0.3 | 0.006 - 0.3 |  |
| c1 (excluded volume) | 1.04 | 1.05 |  |
| c2 (hydration layer) | -2 | -2 |  |
| scale factor | 4.04 | 7.097 |  |
